# Data-Centric Evaluation of Protein Function Prediction Pipelines

**DOI:** 10.64898/2026.08.05.742971

**Authors:** Nicole Soto-García, Norma Murillo-Acevedo, Julián García-Vinuesa, Ana Luisa Islas-Ávila, Mehdi D. Davari, Leandro Murgas-Saavedra, Ahmed Hassanin, Karen Oróstica, Jorge González-Puelma, Marcelo Navarrete, Alicia Martínez-Rebollar, Roberto Uribe-Paredes, Frederic Cadet, David Medina-Ortiz

**Affiliations:** Departamento de Ingeniería en Computación, Universidad de Magallanes, Avenida Bulnes 01855, 6210427, Punta Arenas, Chile; Departamento de Ciencias Computacionales, Tecnológico Nacional de México/CENIDET, Int. Internado Palmira SN, 62490, Morelos, México; Leibniz-Institute of Plant Biochemistry, Department of Bioorganic Chemistry, Weinberg 3, D-06120 Halle, Germany; Department of Pharmacognosy, Faculty of Pharmacy, Assiut University, 71526 Assiut, Egypt; Data Science Institute, Universidad del Desarrollo, Av Plaza 680, 7610615, Santiago, Chile; Centro Asistencial Docente e Investigación, Universidad de Magallanes, Av. Los Flamencos 01364, Punta Arenas, Chile; Escuela de Medicina, Universidad de Magallanes, Avenida Bulnes 01855, Punta Arenas, Chile; PEACCEL, AI for Biologics, Paris, France

**Keywords:** Protein machine learning benchmarking, Data-centric machine learning, Antioxidant proteins, Redundancy reduction, Dataset splitting, Methodological decisions

## Abstract

Performance estimates in protein function prediction depend not only on model choice but also on upstream decisions that define the learning problem. Using antioxidant protein classification as a controlled case study, we evaluated how dataset harmonisation, protein representation, redundancy control, and partitioning strategy affect protein machine learning pipelines. **W**e integrated 18,804 records from 12 publicly available dataset entries into a curated consensus dataset of 4,193 protein sequences. One-hot encoding and six pretrained protein language model representations were evaluated as model inputs and as similarity spaces for redundancy reduction and distance-aware splitting. Representation choice substantially altered dataset geometry, retained dataset size, class balance, and downstream evaluation. At representation-specific p90 thresholds, one-hot encoding retained the complete dataset, whereas pretrained embeddings retained between 5% and 25% of sequences. Distance-aware partitioning reduced apparent performance relative to random splitting by up to 0.15 MCC before redundancy control, while this difference narrowed after similarity filtering. Selected configurations nevertheless maintained high performance under stricter evaluation, reaching an MCC of 0.84. These findings show that performance estimates should be interpreted as outcomes of complete data-centric workflows rather than isolated properties of predictive models.

## 1 Introduction

Machine learning (ML) has become central to computational protein and peptide science, supporting predictive modelling in protein function prediction (Wang et al., 2024; Chen et al., 2025), classification of peptide activity (Zhou et al., 2026; Priyadarsinee et al., 2026; Ibisanmi et al., 2026), and therapeutic discovery (Ekambaram and Dokholyan, 2025; Liu et al., 2025). In protein sequence analysis, recent advances in representation learning, particularly through protein language models (pLM), have improved the extraction of informative sequence-level patterns and expanded the use of computational models for hypothesis generation, candidate prioritisation, and large-scale functional annotation (Hornback et al., 2026; Chhibbar and Das, 2025).

However, predictive performance in biomolecular ML is not determined by model architecture or algorithm choice alone. It is also shaped by upstream decisions that define the learning problem before model training begins, including source selection, dataset integration, label harmonisation, quality control, redundancy reduction, negative-sample construction, representation choice, and train–test partitioning (Bernett et al., 2024; Kapoor and Narayanan, 2023; Heil et al., 2021). We refer to this perspective as data-centric, in which the composition, curation, and structuring of the dataset are treated as first-class determinants of model performance, alongside algorithm choice (Zha et al., 2025; Medina-Ortiz et al., 2026c). These decisions are especially important in protein sequence classification, where datasets often contain homologous, duplicated, or highly similar sequences that can be distributed across training and evaluation partitions. When this structure is not explicitly controlled, models may exploit sequence similarity rather than learn transferable biological patterns, producing overly optimistic estimates of generalisation performance (Bushuiev et al., 2024; Graber et al., 2025; Herrera-Rocha et al., 2025). Importantly, this risk is not limited to sequence-level redundancy, since sequences that differ substantially can still fold into highly similar structures, meaning that sequence-based similarity measures alone may not capture all relevant sources of train–test relatedness.

Dataset construction is therefore not a neutral preprocessing step, but a central component of experimental design. The integration of multiple public sources can modify class balance, annotation confidence, sequence diversity, and the definition of the negative class, all of which influence the apparent difficulty of the prediction task (Aguilera-Puga and Plisson, 2024; Martinez et al., 2025; Fernández-Díaz et al., 2024; Zhang et al., 2026). Similarly, redundancy-control strategies, and partitioning schemes determine how much similarity is allowed between training and test sequences, directly affecting whether reported metrics reflect memorisation of related examples or performance under more stringent evaluation conditions.

Protein representations introduce an additional layer of methodological complexity. In many workflows, representations are used as model-input features, but they are also increasingly used to define similarity relationships among sequences for clustering, redundancy reduction, nearest-neighbour analysis, and distance-aware splitting (Hu et al., 2023; Schütze et al., 2022). This dual role means that a representation does not merely describe each sequence for a classifier. It also defines the geometric space in which sequences are considered close, redundant, or sufficiently separated for evaluation. Different protein representations may therefore induce distinct similarity landscapes over the same dataset and lead to different retained subsets, train–test similarity structures, and performance estimates (Dickson and Mofrad, 2024; Cheng et al., 2026; Pereira et al., 2025; Gujral et al., 2025). These representation-induced similarity landscapes also affect how sequences are assigned to training, validation, and test sets. Under random splitting, closely related or homologous sequences may appear in different partitions, causing validation or test sequences to remain highly similar to training sequences. If this similarity structure is not controlled, evaluation can be affected by data leakage, leading to overestimated predictive performance (Teufel et al., 2023; Joeres et al., 2025).

Recent benchmarking resources and modelling frameworks have addressed specific aspects of this problem. In peptide and protein prediction, PepBenchmark, the Peptide Property Benchmark (PPB), and AutoPeptideML have highlighted how dataset curation, redundancy reduction, homology-aware partitioning, negative-sample definition, and evaluation protocol design can substantially alter reported performance (Zhang et al., 2026; Xiaoying et al., 2026; Fernández-Díaz et al., 2024). Complementary resources for protein sequence and structure learning, including DeepProtein and ProteinShake, have contributed with standardised datasets, benchmark tasks, model libraries, and evaluation procedures for assessing protein representations and predictive models (Xie et al., 2025; Kucera et al., 2023). These studies have improved methodological awareness and reporting standards, but most sources of variability are still examined separately, often across different datasets, representations, and evaluation settings. Consequently, it remains difficult to determine how these decisions interact within a single controlled workflow.

In this work, we perform a data-centric evaluation of protein function prediction pipelines and treat predictive performance as an outcome of interconnected methodological decisions rather than as an isolated property of the supervised learning algorithm. Using antioxidant protein classification as a controlled case study, we assess how dataset harmonisation, protein representation, redundancy control, and distance-aware partitioning reshape dataset composition, similarity structure, train–test separation, and apparent predictive performance across multiple classifiers. The objective is not to construct the best antioxidant protein predictor or to define a universal benchmark, but to examine how upstream methodological decisions modify the interpretation of protein sequence classification performance. By studying protein representations both as model-input features and as similarity spaces for redundancy control and distance-aware splitting, this work provides a reproducible framework for interpreting performance estimates in relation to the methodological choices that generate them.

## 2 Methods

### 2.1 Study design and experimental factors

This study was designed as a data-centric evaluation of machine learning workflows for protein sequence classification (Zha et al., 2025). The objective was to assess how upstream methodological decisions shape the construction of the learning problem, the similarity relationships among sequences, the evaluation scenario, and the resulting interpretation of predictive performance. Antioxidant protein classification was used as a controlled case study because it combines several challenges commonly encountered in biomolecular machine learning, including multi-source data integration, heterogeneous annotations, sequence redundancy, class imbalance, and multiple alternative protein representations (Meng et al., 2023; Lin et al., 2024). Its comparatively small dataset size makes it a computationally tractable test bed for evaluating upstream methodological decisions.

The analytical workflow was organised into four conceptual stages, including data construction, representation and similarity, data structuring, and prediction and interpretation (Figure **1**). The data construction stage included source collection and dataset harmonisation, defining which sequences were retained, how labels were assigned, and how conflicts or low-quality records were handled (Medina-Ortiz et al., 2026c). The representation and similarity stage converted protein sequences into numerical feature spaces and quantified pairwise relationships among sequences (Shaw et al., 2025). The data structuring stage used these similarity relationships to perform redundancy control and to generate random, stratified, and distance-aware train–test partitions (Joeres et al., 2025). The final stage trained supervised machine learning models and interpreted their performance in relation to the methodological decisions that generated each evaluation scenario.

**Figure 1:**
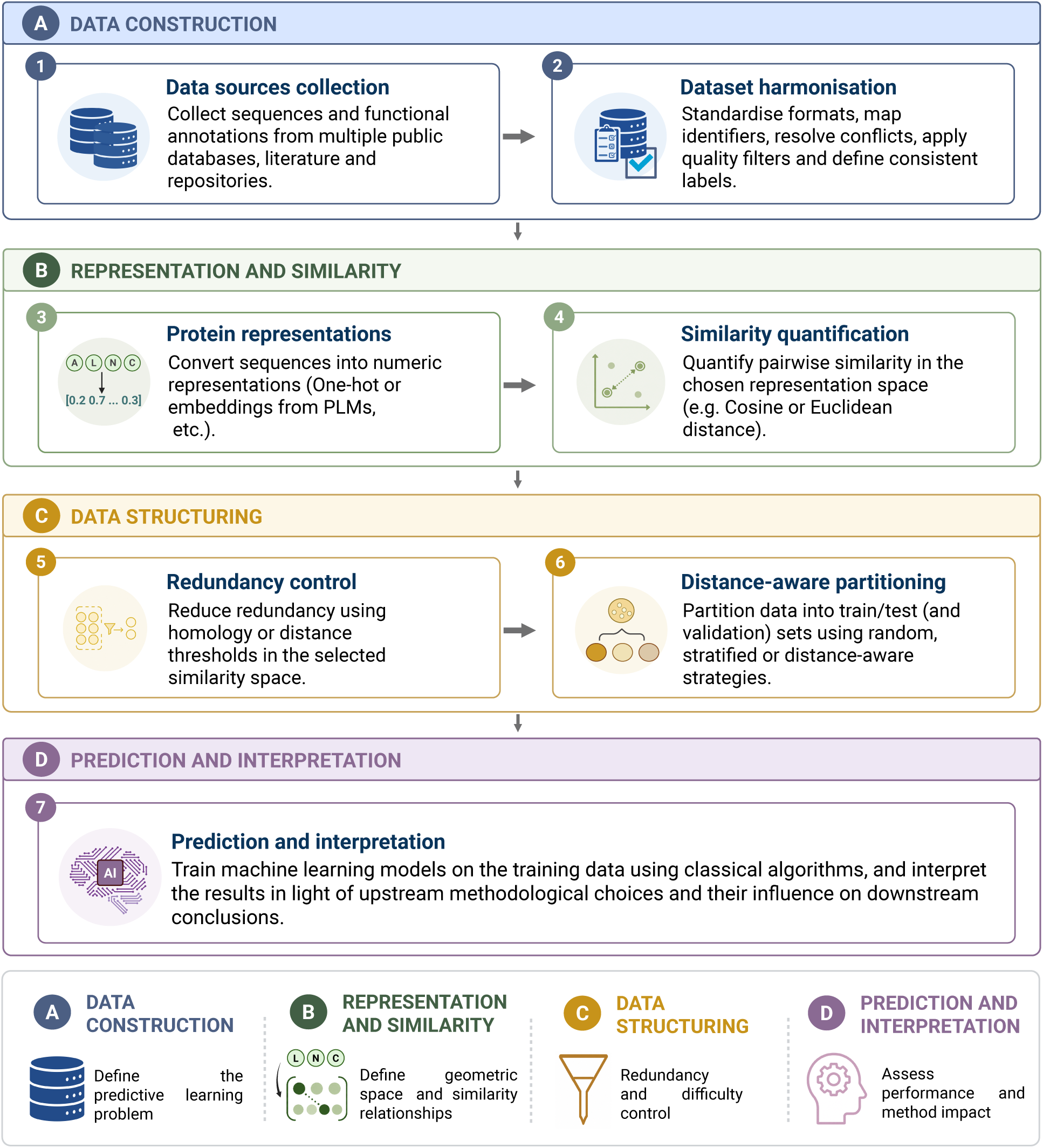
Data-centric workflow for protein function prediction evaluation. The workflow organises protein machine learning evaluation into four interconnected methodological stages. Data construction defines the learning problem through source collection and dataset harmonisation. Representation and similarity define the numerical and geometric spaces in which sequences are encoded and compared. Data structuring controls redundancy and train–test similarity through homology-based or representation-aware filtering and distance-aware splitting. Prediction and interpretation assess model performance while considering how upstream methodological choices shape the conclusions drawn from the evaluation.

The work focused on four methodological dimensions, including dataset construction and harmonisation, protein sequence representation, redundancy-control strategy, and distance-aware partitioning. These dimensions were evaluated through matched supervised learning comparisons whenever possible, so that the effect of a given methodological factor could be interpreted while keeping the remaining workflow components fixed. This separation allowed us to evaluate whether protein representations affect predictive performance only through the features supplied to the model, or also through their role in defining redundancy, train–test separation, and evaluation difficulty.

The main experimental factors are summarised in Table **1**. Dataset-level analyses evaluated the effect of source support and consensus-based harmonisation. Representation-level analyses compared one-hot encoding with pretrained protein language model embeddings. Redundancy-control analyses compared unreduced datasets, homology-based filtering, and representation-distance-based filtering. Partitioning analyses compared random, stratified, and distance-aware splitting strategies. Finally, supervised models were evaluated across repeated random seeds using classification metrics suitable for imbalanced datasets, with Matthews correlation coefficient (MCC) used as the primary performance metric and F1-score as a complementary metric.

**Table 1:**
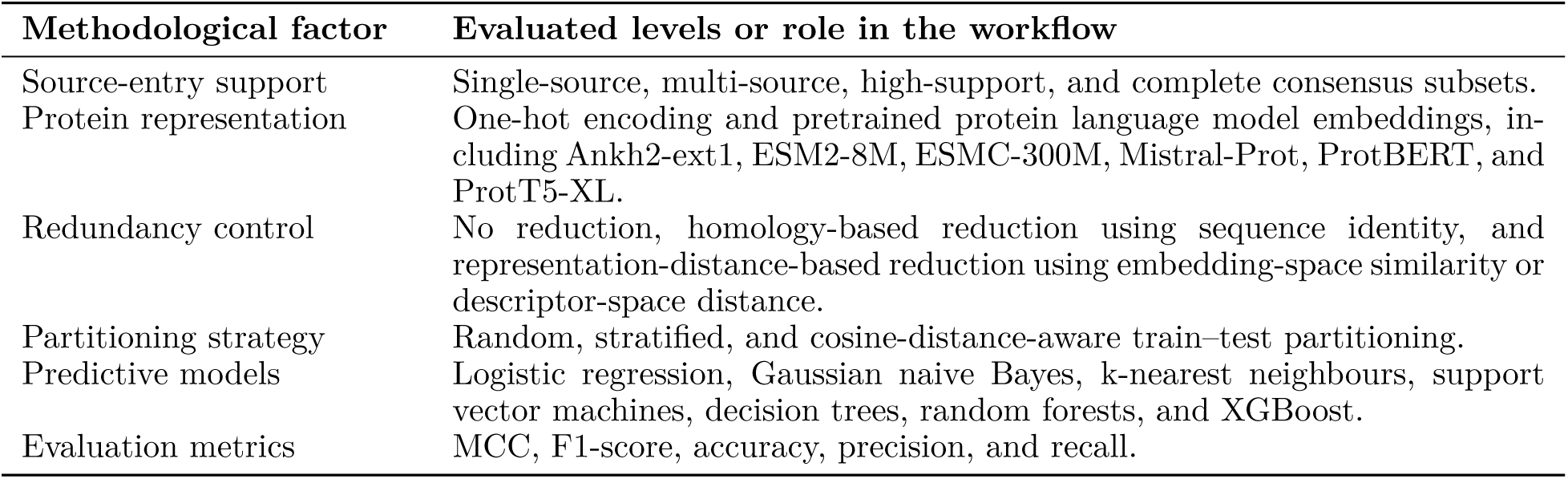
Methodological factors evaluated across the data-centric protein function prediction workflow.

This design treats predictive performance as a property of the complete evaluation workflow, not as an isolated property of the supervised learning algorithm (Bernett et al., 2024; Zha et al., 2025). The purpose was not to identify the best antioxidant protein predictor or to establish a universal benchmark, but to quantify how common data-centric decisions alter dataset composition, similarity structure, evaluation difficulty, and apparent model performance within a reproducible protein function prediction workflow (Walsh et al., 2021). The Supplementary Information (SI) provides detailed descriptions of dataset construction, representation extraction, redundancy-control procedures, splitting strategies, train–test similarity diagnostics, model training and evaluation, paired methodological delta analyses, sensitivity analyses, and implementation details, organised across Sections S1–S8.

### 2.2 Dataset construction and harmonisation

Candidate source-level dataset entries were identified broadly from publicly available publications, databases, benchmark collections, supplementary materials, and repositories associated with antioxidant protein or peptide prediction. Retrieved sources could contain proteins, peptides, precursor sequences, or mixed sequence collections. Source-level dataset entries were retained in the acquisition provenance when their sequence records could be mapped to an antioxidant/non-antioxidant classification context and evaluated using the predefined record-level eligibility criteria. Eligibility for inclusion in the analytical dataset was therefore determined at the record level after source-specific parsing and label standardisation. Sources without accessible sequence-level records were excluded from processing, whereas individual records with ambiguous or non-interpretable labels, malformed sequences, or other unresolved inconsistencies were excluded during source-specific processing.

Each retained source-level dataset entry was processed using a source-specific parsing workflow adapted to the structure of the original files. Input formats included tabular files, FASTA-like records, spreadsheets, supplementary datasets, and dataset partitions distributed as training, testing, positive, negative, or independent subsets. When a publication, database, or repository distributed its data across multiple files or partitions, all associated files were treated as components of the same source-level dataset entry to preserve provenance and enable intra-source consistency checks. Source-level dataset entries containing peptide-level or mixed sequence collections were retained in the acquisition provenance even when some or all of their records were subsequently excluded by the analytical eligibility criteria.

Parsed records were converted into standardised sequence-label tables containing, at minimum, the sequence, the source-level label, and the source identifier. Labels were assigned according to the original source definition, with positive labels corresponding to records reported as antioxidant and negative labels corresponding to records reported as non-antioxidant. No labels were reassigned based on sequence similarity, external functional assumptions, or downstream model behaviour. Source-level dataset entries containing only antioxidant-labelled records were retained as positive-only evidence entries when their labels were explicit, whereas entries containing both classes contributed positive and negative evidence according to their original definitions.

The standardised source-level tables were concatenated and reshaped into a sequence-centred evidence matrix. In this matrix, each row represented a unique sequence and each source-level dataset entry was represented as a separate column containing the label assigned by that entry. Missing source–sequence combinations were treated as absence of evidence and were not interpreted as negative labels. For each sequence, the number of reporting source-level dataset entries, the number of antioxidant assignments, the number of non-antioxidant assignments, and the corresponding antioxidant-label proportion were computed to summarise cross-source evidence.

Consensus labels were assigned from this evidence matrix using strict agreement rules. Sequences reported only as antioxidant across all source-level dataset entries in which they appeared were retained as consensus-positive examples, whereas sequences reported only as non-antioxidant were retained as consensus-negative examples. Identical sequences with concordant labels were collapsed into single entries while preserving source-level evidence. Sequences with mixed evidence, defined as at least one antioxidant and at least one non-antioxidant assignment across source-level dataset entries, were flagged as annotation conflicts, exported for traceability, and excluded from the curated consensus binary dataset. Source-entry support was defined as the number of source-level dataset entries reporting a concordant label for a retained sequence. It was interpreted as documentary concordance rather than as independent experimental validation or a direct measure of label reliability.

Final record-level quality control was applied after consensus assignment. Sequences were retained in the analytical dataset only when they contained canonical amino acids and fell within the accepted length range of 70–1024 residues. The minimum length was used to define the protein-level prediction task and exclude short peptide-like records, whereas the upper bound was determined by the maximum sequence length supported by some of the selected protein language models (Tran et al., 2023). Consequently, a source could remain documented in the acquisition provenance even when none of its retrieved records satisfied the final protein-level eligibility criteria. The same quality-control rules were applied to the consensus-labelled dataset and the conflict file to maintain a consistent audit trail. The resulting curated consensus dataset was used as input for representation extraction, redundancy control, dataset partitioning, and supervised model evaluation. Complete source descriptions, retrieved-file information, parsing rules, evidence-matrix construction, consensus criteria, filtering procedures, and curation summaries are provided in SI Section S1.

### 2.3 Protein sequence representations

The protein sequences in the final curated consensus dataset were transformed into numerical representations prior to redundancy control, dataset partitioning, and supervised model evaluation. To ensure that down-stream comparisons reflected differences in representation strategy rather than differences in input data, the same curated sequence set was used for all encoding procedures.

The study included one sequence-derived baseline and six pretrained protein language model representations. The baseline corresponded to one-hot encoding over the standard amino acid alphabet, with sequences represented as fixed-length residue-level binary matrices (Jing et al., 2019). To obtain vectors of comparable size across proteins, one-hot matrices were zero-padded according to the longest accepted sequence length after dataset curation. Pretrained protein language model embeddings were generated using Ankh2-ext1, ESM2-8M, ESMC-300M, Mistral-Prot, ProtBERT, and ProtT5-XL (Elnaggar et al., 2023; Lin et al., 2022; Candido et al., 2026; Mourad, 2024; Elnaggar et al., 2021). For each protein language model, residue-level embeddings were extracted from the final hidden layer and aggregated into fixed-length sequence-level vectors using mean pooling across residues.

Representations were used in two distinct methodological roles. First, they served as model-input features for supervised classification. Second, they defined numerical similarity spaces used for representation-distance-based redundancy control, train–test similarity analysis, and distance-aware partitioning. These roles were treated independently whenever possible, allowing the representation used for model training to differ from the representation used to define redundancy or distance-aware splitting. This design made it possible to distinguish the predictive effect of a representation from its effect on dataset geometry and evaluation difficulty.

All representations were generated using Sylphy v0.2.0 (Medina-Ortiz et al., 2026a), and sequence identifiers were retained to preserve the correspondence between numerical vectors, original protein sequences, labels, redundancy-control outputs, and partitioning files. Additional details on model identifiers, vector dimensionalities, extraction settings, pooling procedures, output formats, and representation metadata are provided in SI Section S2.

### 2.4 Redundancy control

Redundancy control was applied before dataset partitioning and model evaluation to generate datasets with different levels of sequence similarity (Walsh et al., 2016). Two complementary reduction strategies were evaluated. The first strategy used homology-based filtering, in which protein sequences were clustered according to amino acid sequence identity using MMseqs2 v18.8cc5c through the BioSieve v0.1.0 workflow (Steinegger and Söding, 2017; Medina-Ortiz et al., 2026b). Representative sequences were retained from each cluster at four identity thresholds, corresponding to 90%, 70%, 50%, and 30% sequence identity. Lower identity thresholds imposed stricter homology control by grouping sequences at lower levels of sequence similarity.

The second strategy used representation-distance-based filtering. In this case, redundancy was defined according to the numerical space induced by each protein representation. For pretrained protein language model embeddings, pairwise relationships were quantified using cosine similarity. For the one-hot baseline, distances were quantified using Euclidean distance. Representation-specific percentile thresholds were derived from the empirical similarity or distance distributions and used to define reduction levels within each representation space. The evaluated thresholds corresponded to p30, p40, p50, p60, p70, p80, p90, p95, p97, p98, p99, p99.5, and p99.9. Lower percentile thresholds imposed stronger redundancy control, whereas higher percentiles retained larger fractions of the dataset. The absolute cosine-similarity and Euclidean-distance thresholds corresponding to each percentile and representation are reported in Supplementary Table S11.

The representation used to perform redundancy control was treated as an independent methodological factor. Therefore, the representation space used to identify redundant sequences could differ from the representation later used as input to the supervised classifier. This design was intended to separate the predictive effect of a representation from its effect on dataset geometry and evaluation difficulty.

For each reduction configuration, one representative sequence was retained per redundancy group, and the corresponding reduced dataset, sequence-to-representative mapping, retained and removed sequence summaries, and label distributions were recorded. Additional details on redundancy-reduction settings, threshold definitions, and reduction outcomes are provided in SI Section S3.

### 2.5 Dataset partitioning and train–test similarity

After dataset curation, representation extraction, and redundancy control, each dataset version was partitioned into training, validation, and test subsets. Three partitioning strategies were assessed to generate evaluation scenarios with different assumptions about class balance and sequence similarity. Random partitioning assigned sequences to folds without explicit class-balance or similarity constraints. Stratified partitioning preserved the relative distribution of antioxidant and non-antioxidant labels across subsets whenever possible. The third strategy was cosine-distance-aware partitioning, hereafter referred to as distance-aware partitioning. It used pairwise distances computed from protein representations to reduce proximity between training and test sequences and generate more stringent evaluation scenarios (Joeres et al., 2025).

Partitioning was performed using BioSieve v0.1.0 with five-fold splitting, with one fold held out as the test set in each split (Medina-Ortiz et al., 2026b). This produced an 80/20 train–test division, after which 10% of the training portion was reserved as a validation set for model selection and intermediate evaluation. All partitioning strategies were repeated across 30 independent random seeds. Sequence identifiers were tracked throughout the partitioning process to verify that no identical sequence appeared in more than one subset within the same split.

In the main representation-specific workflow, representation roles were matched whenever possible. When representation-dependent steps were applicable, the same numerical representation was used as model input, as the representation-distance-based redundancy-control space, and as the distance-aware partitioning space. In practice, this means that whichever representation trained the classifier also served to identify redundant sequences and defined train–test proximity during splitting. Thus, each predictive representation was evaluated under a matched dataset-structuring scenario in which redundancy control and distance-aware splitting were defined in the same representation space used for supervised learning. Homology-based reduction was treated separately because it was defined from amino acid sequence identity rather than from representation-space distance. Configurations in which the predictive representation, redundancy-control space, or split space differed were analysed separately as representation-role sensitivity analyses.

Each generated split was checked before model training. Splits were excluded when required train, validation, or test files were missing, when overlap among subsets was detected, or when the resulting class distribution did not allow binary model training or evaluation (only positive or only negative sequences in at least one subset). Invalid splits were recorded by the workflow and excluded from downstream comparisons without interrupting the full experimental run.

Train–test similarity was computed as a post-partitioning diagnostic. For each test sequence, the maximum similarity to any sequence in the corresponding training set was calculated in the matched representation space associated with the evaluated configuration. For random and stratified partitions, this diagnostic was computed *post hoc* using the corresponding predictive representation, allowing direct comparison with distance-aware partitions generated in the same representation space. Split-level summaries included the mean and median maximum train–test similarity, upper-percentile similarity values, the maximum observed similarity, and the proportion of test sequences exceeding predefined similarity cutoffs. These diagnostics were used to verify whether distance-aware partitioning produced more restrictive evaluation scenarios than random or stratified splitting and to support the interpretation of downstream performance differences.

Additional details on partitioning settings, split-validity criteria, representation-space diagnostics, and train– test similarity summaries are provided in SI Section S4.

### 2.6 Model training and evaluation

Supervised learning models were trained to evaluate how methodological decisions affected predictive performance across the generated dataset configurations. The goal of the modelling stage was not to optimise a single antioxidant protein predictor, but to provide a consistent evaluation layer across representations, redundancy-control strategies, and partitioning scenarios.

Seven classical machine learning algorithms were evaluated, covering different modelling paradigms. These included logistic regression, Gaussian naive Bayes, k-nearest neighbours, support vector machines, decision trees, random forests, and XGBoost (Greener et al., 2022). For each algorithm, predefined hyperparameter grids were used to explore comparable model configurations across experimental settings. The complete hyperparameter configurations evaluated for all seven algorithms are reported in Supplementary Table S14.

For each valid split, models were trained using the numerical representation assigned to the corresponding configuration. Model-input features were matched to the active representation under evaluation, whereas redundancy control and partitioning followed the dataset-structuring configuration defined upstream. This ensured that supervised learning was evaluated under the same dataset composition and train–test separation conditions established by the preceding methodological steps. Predictive performance was assessed on the held-out test subset using MCC, F1-score, accuracy, precision, and recall (Tharwat, 2021). Because the curated antioxidant protein dataset was class-imbalanced, MCC was used as the primary metric for comparing configurations (Luque et al., 2019). F1-score was retained as a complementary metric, whereas accuracy, precision, and recall were treated as secondary metrics supporting performance interpretation.

Model evaluations were planned across 30 independent random seeds. Performance summaries were computed across valid repetitions using descriptive statistics, allowing comparison of both average performance and variability across data splits. Invalid configurations, including those lacking sufficient class diversity for model training or test-set evaluation, were excluded from metric aggregation and recorded for reproducibility. Complete model implementations, hyperparameter grids, metric definitions, and aggregation procedures are provided in SI Section S5.

### 2.7 Methodological delta analyses

Methodological delta analyses were used to quantify how specific workflow decisions changed predictive performance relative to matched reference configurations. For a given metric *M*, the delta associated with a candidate configuration *c* and a matched reference configuration *r* was defined as

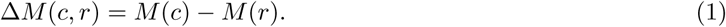

Here, *M* (*c*) and *M* (*r*) denote the performance values obtained under the candidate and reference configurations, respectively. Candidate and reference configurations were matched by dataset composition or redundancy-control condition, as well as by training representation, classifier, random seed, and evaluation metric whenever possible. This matching scheme was used so that the estimated delta primarily reflected the methodological factor under analysis.

For partitioning analyses, random splitting was used as the matched reference. The effect of a partitioning strategy *s* was computed as

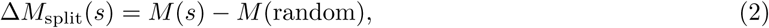

where *s* corresponded to either stratified or distance-aware splitting under the same representation, redundancy-control condition, classifier, and seed. For redundancy-control analyses, unreduced datasets were used as the matched reference. The effect of a reduction condition *q* was computed as

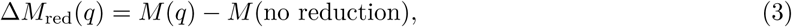

where *q* corresponded to either homology-based reduction or representation-distance-based reduction under matched modelling and partitioning conditions. For representation-role analyses, selected configurations were compared by fixing the predictive representation and varying the representation space used for redundancy control or distance-aware partitioning.

Delta values were computed only for valid matched pairs available after upstream model training. A pair was considered valid when both the candidate and reference configurations had completed model fitting and metric calculation under the corresponding experimental conditions. Invalid or unmatched configurations were recorded but excluded from paired delta summaries.

For each methodological comparison, deltas were summarised across random seeds and valid matched settings. The average delta was calculated as

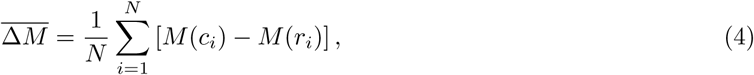

where *N* is the number of valid matched candidate–reference pairs. Positive delta values indicated higher performance under the candidate methodological condition, whereas negative values indicated lower performance relative to the matched reference. These analyses were used to interpret changes associated with splitting strategy, redundancy-control regime, reduction space, and representation role. For paired ranking summaries, the candidate-minus-reference delta was expressed equivalently as MCC loss:

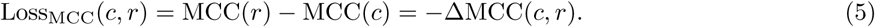

Lower MCC loss values indicated better preservation of, or improvement over, the matched reference performance. Configurations were ranked primarily by MCC loss, with F1-score used as a complementary ranking metric. Accuracy, precision, and recall were treated as secondary metrics.

Complete metric definitions, candidate–reference matching rules, validity criteria, and aggregation procedures for the paired methodological delta analyses are provided in SI Section S6.

### 2.8 Sensitivity analyses

Sensitivity analyses were performed to evaluate whether the main conclusions were consistent across selected dataset-support, similarity-control, and representation-role scenarios. These analyses were designed as targeted methodological checks rather than as an exhaustive benchmark of all possible combinations of representations, redundancy-control strategies, partitioning schemes, classifiers, and thresholds.

The source-support analysis evaluated whether the number of source-level dataset entries reporting concordant labels for each retained sequence influenced dataset composition and downstream predictive performance. Sequence-level subsets were constructed from the final harmonised dataset using this count. Four subsets were defined: single-source sequences reported by exactly one source-level dataset entry, multi-source sequences reported by two or more entries, high-support sequences reported by three or more entries, and the complete consensus dataset, which included all sequences retained after upstream harmonisation, conflict resolution, and filtering. Source-entry support was interpreted as documentary concordance rather than as independent experimental validation. The complete consensus dataset corresponds to the full curated consensus dataset used in the main analyses.

For each source-support subset, dataset size and class balance were summarised before model training. Predictive performance was then evaluated using a common supervised-learning workflow with selected protein representations, partitioning strategies, random seeds, and classifiers.

A redundancy and similarity degradation analysis was used to examine how predictive performance changed as similarity-control constraints became more restrictive. Homology-based degradation compared the unreduced dataset with datasets reduced at 90%, 70%, 50%, and 30% sequence identity. Representation-distance-based degradation compared unreduced datasets with datasets reduced using selected percentile thresholds, including p99, p95, p90, p80, and p70, across selected embedding spaces. These analyses were used to determine whether decreasing redundancy and train–test similarity modified apparent performance and evaluation difficulty.

Finally, representation-role sensitivity analyses were performed to evaluate whether the representation used to define dataset structure influenced performance beyond its role as model input. These analyses were based on targeted non-matched representation-role configurations rather than on a full factorial crossing of all possible training, reduction, and splitting spaces. In the first design, the predictive representation and split space were kept fixed, while the representation-distance-based redundancy-control space was varied. This allowed models trained with the same representation to be compared across datasets reduced using alternative embedding spaces. In the second design, performed on a smaller subset of configurations, the redundancy-control and distance-aware split spaces were kept fixed to a common representation space, while the predictive representation was varied among selected model-input representations. Together, these analyses tested whether the similarity space used for redundancy control, and in selected cases for distance-aware partitioning, could alter reduced dataset composition, train–test similarity, and apparent performance independently of the representation supplied to the classifier.

These sensitivity analyses supported the interpretation of the main workflow by testing whether source-support criteria, similarity-control strength, and representation role altered dataset composition, evaluation difficulty, and model performance. Additional methodological details, including source-support subset definitions, threshold settings for homology-based and representation-distance-based degradation, representation-role configuration summaries, and paired-comparison criteria, are provided in SI Section S7.

### 2.9 Implementation strategy, software, and reproducibility

The computational workflow was implemented as a modular and reproducible analysis pipeline covering dataset harmonisation, representation extraction, redundancy control, dataset partitioning, supervised model training, metric aggregation, and downstream methodological analyses. Each stage was organised with explicit input and output files, allowing individual steps to be re-run, inspected, and validated independently.

Automated execution was coordinated using Snakemake v9.19.0, which was used to preserve dependencies among analysis steps and standardise file generation across the evaluated representation, reduction, partitioning, classifier, and random-seed configurations. Intermediate outputs, including curated datasets, representation matrices, redundancy mappings, split files, model outputs, metric tables, and diagnostic summaries, were stored as traceable workflow artefacts.

Protein representation extraction was performed using Sylphy v0.2.0 (Medina-Ortiz et al., 2026a) for both one-hot encoding and pretrained protein language model embeddings. Redundancy control and dataset partitioning were implemented using BioSieve v0.1.0 (Medina-Ortiz et al., 2026b). Homology-based redundancy reduction used MMseqs2 v18.8cc5c as the sequence-clustering backend (Steinegger and Söding, 2017), whereas representation-distance-based reduction and distance-aware splitting used the numerical representation matrices generated for each configuration. Supervised model training and evaluation were implemented in Python v3.11.5 using NumPy v2.4.6, pandas v3.0.2, SciPy v1.17.1, scikit-learn v1.8.0, XGBoost v3.2.0, and Matplotlib v3.10.8.

Reproducibility was supported by fixed random seeds, explicit configuration files, persistent sequence identifiers, split-integrity checks, and systematic recording of invalid or incomplete configurations. Split-generation workflows checked for missing train, validation, and test files, subset overlap, empty partitions, and insufficient class diversity before downstream model evaluation. Invalid splits or failed model configurations were recorded and excluded from aggregate summaries without interrupting the complete workflow.

All downstream summaries were generated from saved metric, split-diagnostic, and paired-comparison tables rather than from manual recalculation. The workflow preserved the distinction between the representation used for model training, the representation used for redundancy control, and the representation used for distance-aware splitting, allowing matched methodological comparisons to be reconstructed from stored configuration metadata. Detailed implementation information, software versions, execution settings, reproducibility records, and retained workflow artefacts are provided in SI Section S8. Configuration files, workflow rules, execution parameters, curated datasets, processed outputs, and analysis scripts are provided with the public repository and archived outputs described in the Code and Data Availability Statement.

## 3 Results

### 3.1 Source integration and source support reshape dataset composition and evaluation outcomes

The dataset construction workflow showed that source integration is not a neutral preprocessing step, but a determinant of the learning problem evaluated downstream. Starting from 18,804 source-level records collected from 12 publicly available source-level dataset entries, the harmonisation workflow identified 7,870 unique amino acid sequences after duplicate resolution and cross-entry integration (Figure **2**A). Among these sequences, 43 were excluded because they showed conflicting labels across source-level dataset entries. The remaining consensus-labelled sequences were further filtered using canonical amino acid and length criteria, resulting in a final curated consensus dataset of 4,193 proteins, including 1,010 antioxidant proteins and 3,183 non-antioxidant proteins.

**Figure 2:**
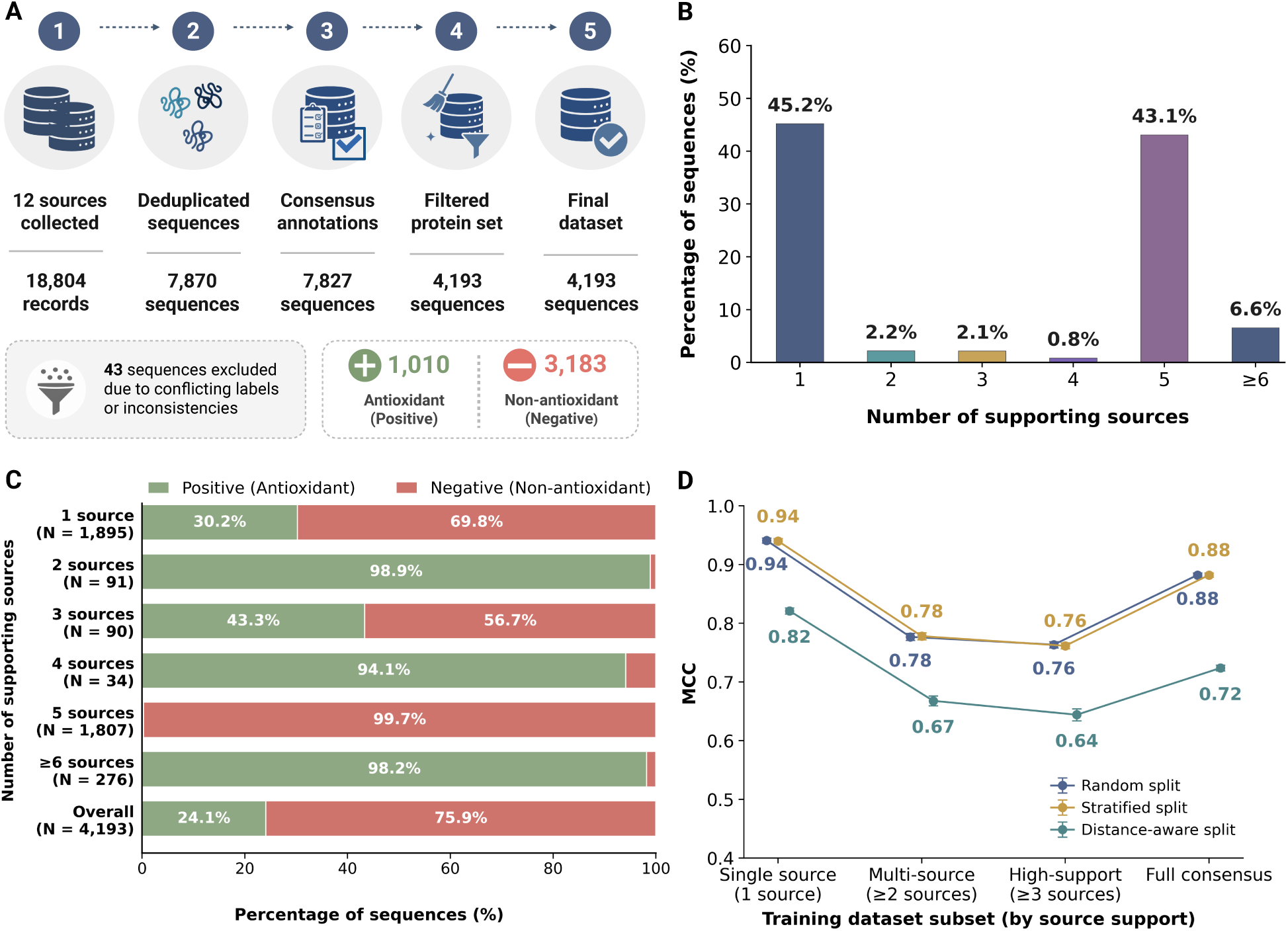
Source support reshapes dataset composition and evaluation outcomes. **(A)** Dataset construction workflow from 18,804 source-level records collected from 12 publicly available source-level dataset entries to the final curated consensus dataset of 4,193 proteins. **(B)** Distribution of sequences according to the number of source-level dataset entries providing concordant support for their final consensus annotation. **(C)** Class composition across source-support categories, showing the percentage of antioxidant and non-antioxidant proteins at each support level. **(D)** Predictive performance across source-support-based subsets under random, stratified, and distance-aware partitioning. Points indicate mean test-set MCC across valid evaluations, and error bars indicate 95% confidence intervals.

Source-entry support was heterogeneously distributed across the final consensus dataset (Figure **2**B). Most sequences were concentrated in the one-entry and five-entry categories. Sequences reported by exactly one source-level dataset entry accounted for 1,895 proteins, corresponding to 45.2% of the final dataset, whereas sequences reported concordantly by five entries accounted for 1,807 proteins, corresponding to 43.1%. The remaining categories represented smaller fractions of the dataset: 91 sequences were reported concordantly by two entries (2.2%), 90 by three entries (2.1%), 34 by four entries (0.8%), and 276 by six or more entries (6.6%). When these categories were aggregated, 2,298 proteins, corresponding to 54.8% of the final dataset, were reported concordantly by two or more source-level dataset entries, and 2,207 proteins, corresponding to 52.6%, were reported concordantly by three or more entries. These counts provide a provenance-aware summary of cross-entry label concordance and describe the annotation structure represented in the final dataset.

The distribution of antioxidant and non-antioxidant labels varied strongly across source-support categories (Figure **2**C). Although the overall positive proportion was 24.1%, this global class balance did not hold uniformly across support levels. Single-source sequences showed a positive proportion of 30.2%, whereas sequences supported by two, four, and six or more sources were strongly dominated by antioxidant proteins, with positive proportions of 98.9%, 94.1%, and 98.2%, respectively. In contrast, the five-source category was almost entirely dominated by non-antioxidant proteins, with only 0.3% positives. The three-source category showed a more balanced composition, with 43.3% antioxidant and 56.7% non-antioxidant proteins. These patterns show that source-entry support reflects not only annotation redundancy and confidence but also shapes the class composition of the dataset, thereby altering the supervised learning problem.

Source-support structure also affected predictive evaluation (Figure **2**D). Across valid representation– algorithm–seed evaluations, random and stratified partitioning produced nearly identical MCC trends across source-support-based training subsets. However, performance did not increase monotonically with source support. Instead, the single-source subset showed the highest MCC under conventional partitioning, with values close to 0.94 for both random and stratified splits, followed by the full consensus dataset with MCC values close to 0.88. In contrast, the multi-source and high-support subsets showed lower MCC values, approximately 0.78 and 0.76, respectively. This non-monotonic pattern indicates that source-entry support should not be interpreted as a simple proxy for increasing classification difficulty, annotation confidence, or predictive performance. Rather, each source-support subset defines a distinct learning problem with its own class balance, sequence composition, and similarity structure.

Distance-aware partitioning produced consistently lower MCC values across all source-support subsets. MCC decreased to approximately 0.82 for the single-source subset, 0.67 for the multi-source subset, 0.64 for the high-support subset, and 0.72 for the full consensus dataset. The consistent reduction relative to random and stratified partitioning indicates that conventional partitions benefit from train–test proximity among related sequences, whereas distance-aware evaluation imposes a more stringent generalisation scenario. Importantly, the decrease was not uniform across source-support subsets, suggesting that the effect of distance-aware partitioning depends on the composition and similarity structure of the subset being evaluated.

These results demonstrate that dataset construction decisions define the evaluation scenario before model training begins. Source integration, conflict removal, consensus labelling, quality control, and source-support filtering jointly determine dataset size, class balance, annotation structure, sequence-similarity structure, and apparent predictive performance. Consequently, model performance in this task should be interpreted in relation to the construction history of the dataset and not only as a property of the supervised learning algorithm.

### 3.2 Protein representations induce distinct similarity landscapes over the same dataset

The final curated consensus dataset was next encoded using one-hot representation and six pretrained protein language model embeddings to evaluate whether the same protein sequences generated comparable or representation-specific similarity structures. This analysis showed that protein representations do not only transform sequences into numerical features for supervised learning. They also define the geometric space in which sequences are considered close, distant, redundant, or separable. Figure **3** summarises these representation-dependent effects using selected embeddings and representative redundancy-control outputs. ESMC-300M and ProtBERT were also included in the complete representation-level analyses, with their similarity profiles, redundancy-reduction outcomes, and distance-degradation results reported in Supplementary Figure S2 and Supplementary Tables S11 and S22.

**Figure 3:**
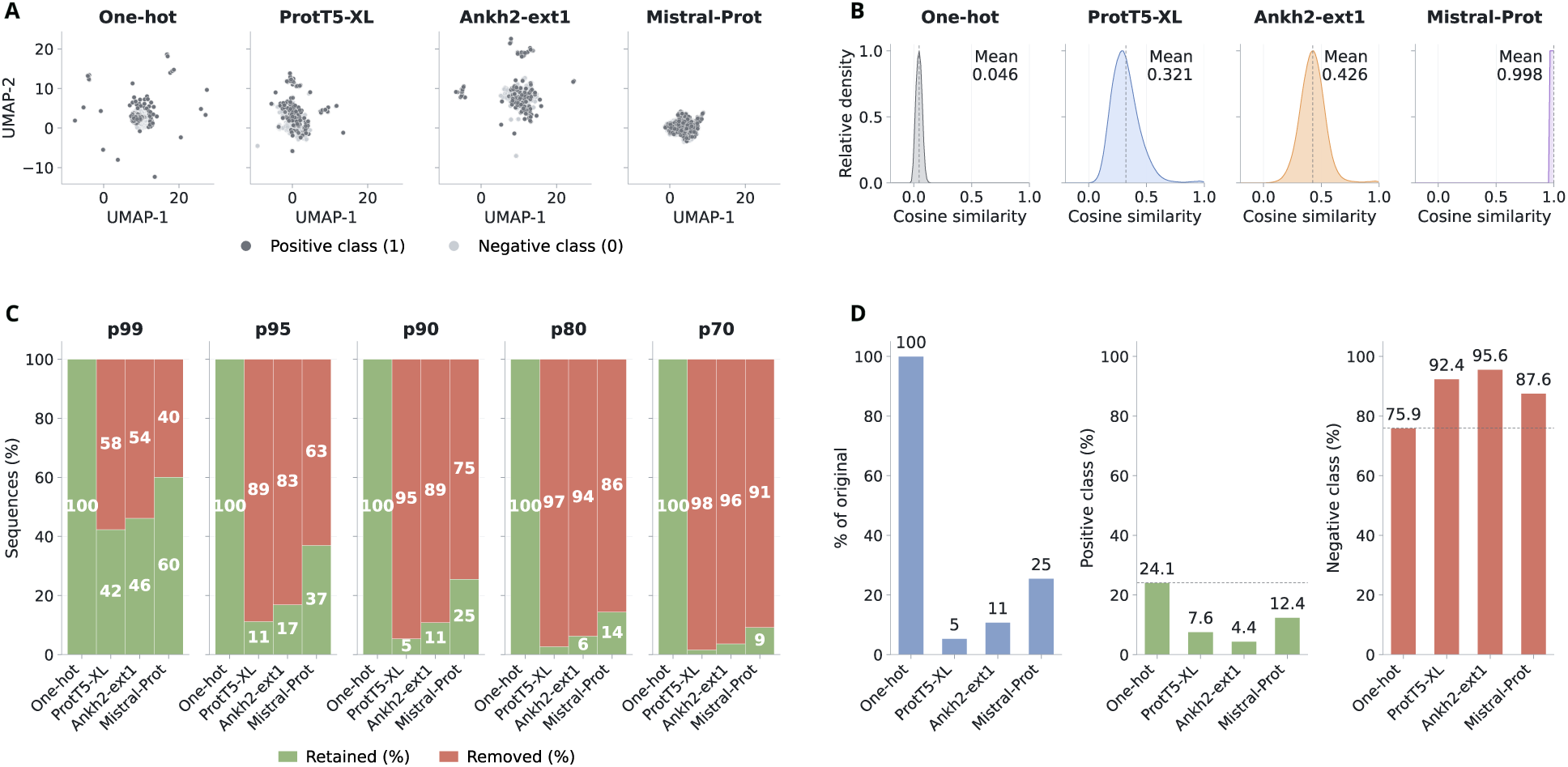
Protein representations induce distinct similarity landscapes and redundancy-control outcomes. **(A)** Two-dimensional UMAP visualisations of sequence representation spaces, showing how the same antioxidant protein dataset is organised under one-hot encoding and representative protein language model embeddings. Points represent proteins and colours indicate antioxidant and non-antioxidant labels. **(B)** Pairwise cosine-similarity distributions across selected representation spaces, with dashed lines indicating the mean cosine similarity for each representation. Redundancy reduction used cosine similarity for pretrained protein language model embeddings and Euclidean distance for one-hot encoding. **(C)** Proportion of retained and removed sequences after representation-distance-based redundancy reduction using selected percentile thresholds. **(D)** Dataset coverage and class-balance changes at the representation-specific p90 thresholds, illustrating how representation-aware filtering can modify both retained dataset size and class composition. Complete results across evaluated representations, thresholds, homology-based reductions, partitioning strategies, and model configurations are provided in the Supplementary Information.

The two-dimensional projections showed that the same dataset adopted markedly different spatial organisations across representation spaces (Figure **3**A). One-hot encoding produced a compact structure with several isolated sequences, reflecting a sparse residue-level encoding dominated by direct sequence composition and positional information. ProtT5-XL and Ankh2-ext1 generated broader embedding spaces with more dispersed local structures, while Mistral-Prot produced a dense and compact distribution. Across all visualised spaces, antioxidant and non-antioxidant proteins were largely intermixed, with no representation producing a clearly separated class-specific geometry. This indicates that the classification task is not defined by an obvious global separation between positive and negative proteins in the reduced representation space.

Pairwise cosine similarity distributions confirmed that each representation induced a distinct similarity landscape over the same set of sequences (Figure **3**B). One-hot encoding produced a narrow distribution centred near zero, with a mean cosine similarity of 0.046, indicating low average similarity between most sequence pairs in this representation. ProtT5-XL and Ankh2-ext1 produced broader distributions centred at intermediate values, with mean cosine similarities of 0.321 and 0.426, respectively. In contrast, Mistral-Prot produced a highly concentrated distribution near one, with a mean cosine similarity of 0.998, indicating that most sequence pairs were placed very close to each other in that embedding space.

These differences show that representation choice changes the effective notion of sequence similarity used by the workflow. A pair of proteins may appear distant in one-hot space, moderately related in ProtT5-XL or Ankh2-ext1 space, and highly similar in Mistral-Prot space. Therefore, representation choice affects not only the information supplied to the classifier, but also the similarity assumptions used during redundancy control, train–test similarity analysis, and distance-aware partitioning.

This result is important for interpreting downstream analyses based on percentile thresholds. Percentile-based criteria are computed within the representation-specific similarity or distance distribution, so the same percentile does not correspond to the same absolute cutoff or the same biological separation across representations. Consequently, redundancy reduction and distance-aware evaluation should be interpreted as representation-dependent operations. The next analysis therefore evaluated how these distinct representation spaces translated into differences in retained dataset size and class composition.

### 3.3 Representation-aware redundancy control modifies dataset coverage and class balance

The representation-dependent similarity landscapes described above produced substantial differences in redundancy-control outcomes. When percentile-based reduction thresholds were applied within each representation space, the proportion of retained sequences varied markedly across embeddings and thresholds (Figure **3**C). This indicates that representation-aware redundancy control does not simply remove duplicated or near-identical sequences. It defines a representation-specific subset of the dataset according to the geometry induced by the selected numerical space.

This pattern was observed across the evaluated percentile thresholds, but p90 was used as an illustrative operating point for detailed reporting because it retained sufficient valid downstream partitions for paired evaluation while still imposing substantial representation-aware redundancy control. At this threshold, one-hot encoding retained 100% of the dataset, indicating that this representation did not identify redundant groups under the evaluated distance criterion. In contrast, pretrained protein language model embeddings produced much stronger reductions. ProtT5-XL retained 5% of sequences, Ankh2-ext1 retained 11%, and Mistral-Prot retained 25%. Thus, the same percentile-based reduction level generated very different dataset sizes depending on the representation used to compute pairwise relationships.

This behaviour was also observed across other percentile thresholds. As expected, lower percentile thresholds imposed stronger redundancy control and retained fewer sequences, whereas higher percentiles were more permissive. However, the magnitude of this effect was representation-dependent. Some embeddings produced rapid reductions even at high percentile thresholds, while others retained larger fractions of the dataset across the same percentile range. These results show that percentile thresholds provide a relative criterion within each representation space, but they do not impose equivalent absolute reduction strength across representations.

Representation-aware redundancy control also modified class balance (Figure **3**D). In the unreduced final curated consensus dataset, antioxidant proteins represented 24.1% of the sequences. At the representation-specific p90 thresholds, the retained positive proportion differed substantially across representation spaces, decreasing to 7.6% for ProtT5-XL, 4.4% for Ankh2-ext1, and 12.4% for Mistral-Prot. The corresponding retained datasets were therefore not only smaller, but also more strongly dominated by non-antioxidant proteins than the original dataset.

These shifts indicate that changes in performance after redundancy reduction cannot be interpreted as the effect of similarity filtering alone. Redundancy control simultaneously changes dataset coverage, class balance, retained sequence composition, and the similarity relationships available for downstream partitioning. Consequently, the representation used for reduction becomes part of the evaluation design itself. Performance estimates obtained after representation-aware filtering must therefore be interpreted in relation to the specific embedding space and threshold used to construct the reduced dataset.

### 3.4 Distance-aware partitioning reveals performance sensitivity while preserving robust configurations

After evaluating how source support, representation geometry, and redundancy control altered dataset composition, we assessed whether these upstream decisions changed the apparent difficulty of model evaluation. We focused on the relationship between train–test similarity, partitioning strategy, redundancy threshold, and predictive performance. Figure **4** summarises this analysis using maximum train–test similarity distributions, matched ΔMCC comparisons, and representation-specific threshold effects.

**Figure 4:**
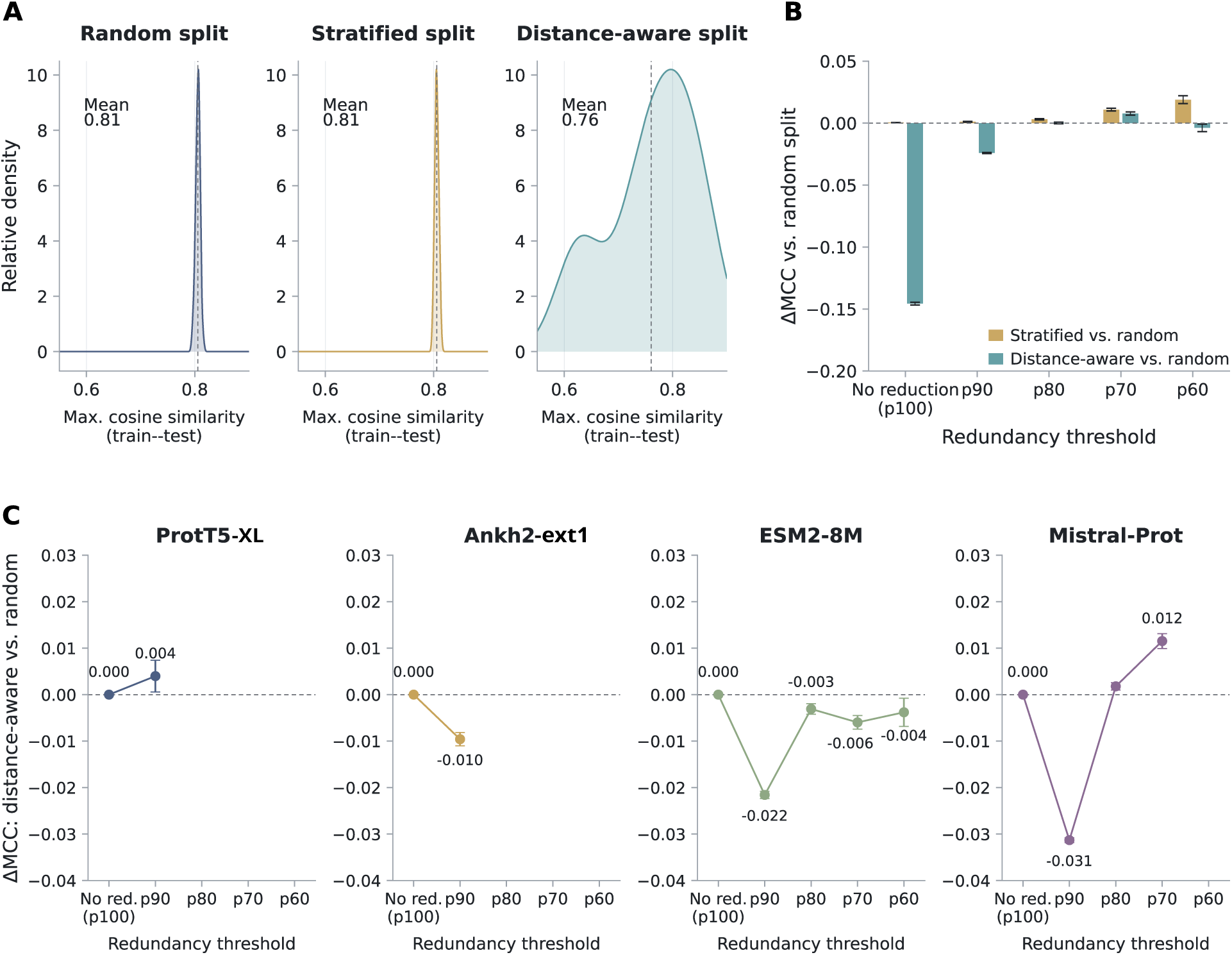
Distance-aware partitioning reveals performance sensitivity to train–test similarity. **(A)** Distribution of maximum train–test cosine similarity under random, stratified, and distance-aware partitioning strategies. **(B)** Change in MCC relative to random partitioning across redundancy thresholds. Bars show ΔMCC for stratified and distance-aware splits relative to the corresponding random-split reference. **(C)** Representation-specific ΔMCC under distance-aware partitioning relative to random partitioning across redundancy thresholds.

Train–test similarity differed substantially across partitioning strategies (Figure **4**A). Random and stratified splits produced highly concentrated maximum-similarity distributions with similar mean values, both centred around 0.81. In contrast, distance-aware partitioning produced a broader distribution and a lower mean maximum train–test similarity of approximately 0.76. This shift indicates that conventional random and stratified partitions preserved highly similar training examples for many test proteins, whereas distance-aware partitioning generated evaluation scenarios with greater separation between training and test subsets.

The performance impact of this separation depended on the redundancy structure of the dataset (Figure **4**B). Stratified partitioning produced ΔMCC values close to zero across the evaluated thresholds, indicating that preserving class balance alone did not substantially change performance relative to random splitting. Distance-aware partitioning produced the strongest decrease in MCC in the unreduced dataset, where train– test similarity was still largely uncontrolled. After representation-distance-based redundancy reduction, the performance gap between random and distance-aware partitioning became smaller and approached zero for several thresholds. This pattern indicates that redundancy control and distance-aware splitting act on related aspects of evaluation difficulty. When redundancy is not explicitly controlled, distance-aware splitting exposes a stronger dependence on train–test proximity. Once redundancy has already been reduced, the additional effect of distance-aware splitting becomes less pronounced.

Representation-specific analyses showed that this effect was not uniform across embedding spaces (Figure **4**C). Some representation and threshold combinations showed negative ΔMCC values under distance-aware splitting, with the largest decreases observed around p90 for selected spaces, including Mistral-Prot and ESM2-8M. In other configurations, the effect was smaller, close to zero, or weakly positive after stronger redundancy control. These non-monotonic patterns indicate that distance-aware evaluation does not impose a fixed performance penalty. Its effect depends on the interaction between the representation used to define similarity, the threshold used to reduce redundancy, the retained class composition, and the final train–test separation achieved by the split.

Not all representation–threshold combinations produced valid evaluation scenarios. Some reduced datasets became too small or too imbalanced to generate training, validation, and test partitions containing both classes. These invalid configurations were excluded from paired delta summaries but retained in the reproducibility record; representative cases and the corresponding numbers of valid and filtered seeds are reported in Supplementary Table S23. This behaviour shows that strong similarity filtering can alter dataset composition enough to compromise binary evaluation. Therefore, threshold selection should be interpreted not only as a redundancy-control choice, but also as a decision that affects class diversity, split validity, and the feasibility of downstream model comparison.

Despite the stricter evaluation imposed by distance-aware partitioning, selected configurations retained high predictive performance in the targeted representation-role sensitivity analysis (Table **2**). In this analysis, three representation roles were considered separately. The model-input representation corresponds to the feature space used to train the supervised classifier. The reduction setting identifies the representation space and redundancy threshold used for representation-distance-based filtering, whereas the split space identifies the representation used to compute train–test proximity during distance-aware partitioning. MCC values were reported as the mean *±* 95% confidence interval computed from valid aggregated test-set MCC values, and only configurations with at least 30 unique matched seeds were retained. The corresponding ΔMCC represents the mean performance difference relative to the matched random-split reference. These configurations are reported descriptively and do not represent the selection of a final antioxidant protein predictor.

**Table 2:** Selected representation-role configurations from the targeted sensitivity analysis under distance-aware partitioning. Configurations are ordered by mean test-set MCC and report the model-input representation, redundancy-reduction setting, split space, algorithm, MCC, and matched ΔMCC.

| Rank | Input rep. | Reduction setting | Split space | Algorithm | MCC mean ± 95% CI | $\Delta\text{MCC}$ |
| --- | --- | --- | --- | --- | --- | --- |
| 1 | ProtT5-XL | Mistral-Prot p90 | Mistral-Prot | SVC cfg. 2 | $0.84 \pm 0.003$ | −0.005 |
| 2 | Ankh2-ext1 | Mistral-Prot p90 | Mistral-Prot | SVC cfg. 2 | $0.82 \pm 0.004$ | −0.02 |
| 3 | Ankh2-ext1 | Mistral-Prot p90 | Mistral-Prot | XGBoost cfg. 3 | $0.79 \pm 0.004$ | −0.008 |
| 4 | Ankh2-ext1 | Mistral-Prot p90 | Mistral-Prot | k-NN cfg. 4 | $0.75 \pm 0.007$ | −0.02 |
| 5 | ProtT5-XL | Mistral-Prot p90 | Mistral-Prot | XGBoost cfg. 3 | $0.74 \pm 0.006$ | −0.03 |

The configuration with the highest observed MCC in this analysis used ProtT5-XL as the model-input representation, Mistral-Prot with a p90 threshold as the redundancy-reduction setting, Mistral-Prot as the distance-aware split space, and an SVC classifier. This configuration reached an MCC of 0.84 *±* 0.003, with a ΔMCC of *−*0.005 relative to its matched random-split reference. Other selected configurations used Ankh2-ext1 or ProtT5-XL as model-input representations with SVC, XGBoost, or k-NN classifiers, reaching MCC values between 0.74 *±* 0.006 and 0.82 *±* 0.004. Across the configurations shown, the reduction setting was consistently Mistral-Prot at p90, and the distance-aware split space was consistently Mistral-Prot, whereas the model-input representation and classifier varied.

These results show that similarity-aware evaluation does not simply lower performance. Instead, it reveals which configurations remain strong after controlling train–test proximity. High MCC values can still be achieved under stricter evaluation, but they depend on the combined configuration of the model-input representation, redundancy-control space, partitioning space, and classifier. Therefore, performance estimates in protein function prediction should be interpreted as properties of complete evaluation workflows rather than as isolated outputs of supervised-learning algorithms.

## 4 Discussion

Using antioxidant protein classification as a controlled case study, we found that predictive performance was shaped not only by the supervised-learning algorithm, but also by the complete evaluation workflow defining the prediction problem. Source integration, consensus labelling, representation choice, redundancy control, and train–test partitioning jointly altered dataset composition, similarity structure, evaluation difficulty, and apparent model performance. These results support a data-centric interpretation of biomolecular machine learning, in which performance estimates are treated as properties of the full experimental design rather than as isolated indicators of model quality.

A first implication is that dataset construction should be considered part of model evaluation. The integration of 12 publicly available source-level dataset entries did not simply increase sample size. It introduced a structured evidence landscape in which source support was unevenly distributed and strongly associated with class composition. More than half of the final dataset was associated with concordant support from multiple source-level dataset entries, but support categories differed markedly in their antioxidant and non-antioxidant proportions. As a result, source-support filtering changed both the annotation structure and the supervised learning problem. The MCC patterns observed across source-support subsets were non-monotonic: single-source and full-consensus datasets showed higher performance under conventional partitioning, whereas multi-source and high-support subsets showed lower performance. This indicates that source support should not be interpreted as a simple proxy for increasing annotation confidence, classification difficulty, or predictive performance. Importantly, distance-aware partitioning reduced MCC across all source-support subsets, indicating that source-support structure and train–test similarity jointly shape the evaluation outcome.

These observations reinforce the need to report dataset provenance, label harmonisation rules, conflict handling, and source-support structure in protein and peptide prediction studies. Public biological datasets are often reused across multiple benchmarks, modified across publications, or distributed as positive, negative, training, testing, and independent subsets with heterogeneous definitions. Without explicit reconstruction of this provenance, models can be evaluated on datasets whose internal evidence structure is poorly understood. In this context, consensus labels and source-entry support should not be treated as passive metadata. They define which examples enter the learning problem, how their labels are documented across source-level dataset entries, and how class balance is shaped before model training begins.

A second implication is that protein representations influence evaluation in more than one way. Protein language model embeddings are commonly interpreted as improved feature spaces for classification, but our results show that they also define the similarity landscape used to identify redundancy, construct distance-aware partitions, and diagnose train–test proximity. The same protein sequences produced sharply different pairwise similarity distributions depending on the representation. One-hot encoding yielded low average cosine similarity, ProtT5-XL and Ankh2-ext1 produced intermediate similarity distributions, and Mistral-Prot placed most sequence pairs close to one another in cosine space. These differences mean that a fixed percentile threshold does not represent an equivalent biological or geometric constraint across representations.

This point is especially important for representation-aware redundancy control. At the representation-specific p90 thresholds, retained dataset size varied from 5% with ProtT5-XL to 25% with Mistral-Prot, while one-hot retained the full dataset under the evaluated criterion. Class balance also changed substantially after reduction, with retained positive proportions decreasing below the original dataset proportion in the protein language model spaces. Therefore, redundancy reduction should not be interpreted only as the removal of duplicate or near-identical sequences. When performed in representation space, it constructs a new dataset whose coverage, class balance, and similarity structure depend on the embedding used to define proximity. The representation is therefore part of the evaluation design itself.

This finding extends previous concerns about data leakage and overoptimistic evaluation in biological machine learning. Studies such as AutoPeptideML have shown that homology-aware partitioning and negative-sample definition can strongly affect reported peptide bioactivity performance (Fernández-Díaz et al., 2024). PepBenchmark similarly emphasised that redundancy removal and biologically informed splitting strategies are essential for reliable peptide benchmarking (Zhang et al., 2026). Our results complement these studies by showing that the embedding space used to define similarity can also determine how much redundancy is removed, how much class imbalance is introduced, and how stringent the evaluation scenario becomes. In other words, leakage control is not only a question of whether redundancy reduction or distance-aware partitioning is applied. It also depends on the space in which similarity is measured.

The interaction between redundancy control and partitioning was also central to the observed performance patterns. Distance-aware partitioning reduced train–test similarity and produced lower MCC than random splitting in the unreduced dataset, consistent with the expectation that conventional random partitions can preserve highly similar training examples for many test sequences. However, after representation-distance-based redundancy reduction, the gap between random and distance-aware splitting narrowed and sometimes approached zero. This suggests that redundancy reduction and distance-aware partitioning act on related sources of evaluation difficulty. When redundancy remains uncontrolled, distance-aware splitting exposes a stronger dependence on train–test proximity. Once similarity has already been reduced upstream, the additional effect of distance-aware splitting becomes smaller and more dependent on the retained dataset structure.

The representation-specific delta analyses further show that stricter evaluation does not impose a uniform penalty. The effect of distance-aware partitioning varied across embedding spaces and thresholds, with non-monotonic patterns across p90, p80, p70, and p60. This behaviour likely reflects the combined influence of similarity distribution, retained subset size, class balance, and split feasibility. Some thresholds produced invalid configurations because the reduced datasets no longer contained sufficient class diversity across train, validation, and test subsets. These invalid cases are not merely technical failures. They reveal that strong filtering can transform the learning problem enough to compromise meaningful binary evaluation. Thus, threshold selection should be reported not only as a redundancy-control parameter, but also as a determinant of downstream evaluation validity.

Despite these constraints, strong performance remained achievable under similarity-aware evaluation. The highest observed MCC was 0.84 *±* 0.003, with only a small decrease relative to the matched random-split reference. This is an important result because the study does not argue that high performance in protein function prediction is necessarily artefactual. Instead, it shows that high performance is more interpretable when it persists under explicit control of source structure, redundancy, and train–test similarity. These outcomes were associated with combinations of model-input representation, reduction space, splitting space, and classifier. This supports a workflow-level interpretation of performance, where robust prediction emerges from the compatibility among dataset construction, representation geometry, similarity control, and modelling strategy.

Several limitations should be considered. First, antioxidant protein classification was used as a controlled case study. Although the methodological principles are relevant to other protein and peptide prediction tasks, the magnitude of each effect will depend on the redundancy, annotation quality, class imbalance, and functional diversity of the target dataset. Second, the representation-role analyses were targeted and not fully factorial across all possible combinations of input representation, reduction space, partitioning space, classifier, and threshold. This choice was necessary to keep the workflow interpretable and computationally tractable, but it leaves higher-order interactions for future work. Third, the study focused on classical supervised classifiers trained on fixed sequence-level representations. Fine-tuned protein language models, deep neural architectures, multimodal protein representations, and structure-aware embeddings may produce different performance profiles and should be evaluated under similar data-centric controls.

Rather than aiming to identify the best-performing model or exhaustively test every possible combination, this study underscores the importance of making the methodological decisions behind protein machine learning pipelines transparent and visible, since each choice shapes the performance ultimately observed. Performance metrics should be reported together with the dataset construction protocol, label harmonisation strategy, redundancy-control method, similarity space, partitioning scheme, random-seed structure, invalid configuration handling, and split-level train–test similarity diagnostics. Without this information, comparisons between models can conflate algorithmic improvement with differences in problem formulation. A model that appears superior under one dataset construction and partitioning regime may not remain superior under a stricter similarity-aware evaluation.

Overall, the results argue for a shift from model-centred benchmarking towards data-centric workflow evaluation. In this view, model performance is the final observable outcome of a sequence of methodological decisions. Each decision shapes the examples available to the classifier, the geometry in which they are compared, the redundancy allowed across partitions, and the difficulty of the final test scenario. Treating these decisions as explicit components of benchmark design may improve reproducibility, facilitate fairer model comparisons, and reduce the risk of overinterpreting performance gains driven by dataset structure. This perspective is especially important as protein language models and large-scale biological datasets become increasingly central to protein function prediction, peptide discovery, and computational protein engineering.

## 5 Conclusion

This work demonstrates that protein function prediction performance is shaped by the complete evaluation workflow, not only by the supervised-learning algorithm. In a controlled antioxidant protein classification case study, source integration, source-support structure, representation geometry, redundancy control, and distance-aware partitioning all modified the dataset being evaluated and the difficulty of the prediction task. Protein representations played a dual role by providing model-input features and defining the similarity spaces used for redundancy reduction and train–test separation.

The main practical conclusion is that performance estimates in biomolecular machine learning should be interpreted together with the methodological decisions that generated them. In the evaluated setting, high predictive performance remained achievable under stricter similarity-aware evaluation, but its interpretation depended on the dataset construction protocol, representation space, redundancy threshold, partitioning strategy, and classifier configuration. Reporting these components explicitly is therefore important for supporting reproducible, transparent, and comparable protein machine-learning studies.

By framing performance as an outcome of data-centric workflow design, this study provides, within the evaluated setting, a reproducible basis for examining protein prediction pipelines beyond model selection alone. Within the scope of this controlled case study, this perspective could support more reliable benchmarking practices and help distinguish predictive generalisation from performance driven by dataset composition, redundancy, or train–test similarity. These conclusions should be interpreted within the modelling setting considered here, which used fixed sequence-level representations and classical supervised classifiers.

Future work should evaluate the transferability of this data-centric framework across additional protein and peptide prediction tasks and alternative sequence- and structure-aware representations. Such evaluations will help distinguish task-specific effects from methodological patterns that recur across biological machine-learning applications.

## Author contributions statement

NS-G, NM-A, and DM-O: conceptualization. NS-G, NM-A, FC, and DM-O: methodology. DM-O: validation. NS-G, NM-A, JG-V, ALI-A, LM, and AH: investigation. NS-G, NM-A, JG-V, ALI-A, KO, JG-P, MN, FC, and DM-O: writing and editing. M.D.D., RU-P, and DM-O: supervision and funding resources. FC and DM-O: project administration. All authors reviewed and approved the final version of the manuscript.

## Code and Data Availability Statement

All source code, workflow implementations, analysis scripts, processed datasets, metadata records, data partitions, model outputs, and other materials required to reproduce the analyses are publicly available in the GitHub repository at https://github.com/kren-ai-lab/protein_ml_decision_analysis. An archived version of the repository and associated research materials is also available through the KREN-AI Lab Zenodo community at https://zenodo.org/communities/kren-ai-lab, under DOI: 10.5281/zenodo.21709987.

## Acknowledgements

NS-G, NM-A, JG-V, and DM-O acknowledge funding from FONDECYT Iniciación 11250295. NS-G acknowledges funding from “Fondos Concursables Postgrado: Fortalecimiento de Tesis y Actividad Formativa Equivalente” (MAG23992). DM-O gratefully acknowledges support from the Centre for Biotechnology and Bioengineering - CeBiB (PIA project FB0001 and AFB240001, ANID, Chile). M.D.D. acknowledges financial support from the German Federal Ministry for Research, Technology and Space (BMFTR) (grant number: 031B1442E). AH is supported by a Ph.D. scholarship from the Ministry of Higher Education of the Arab Republic of Egypt. ALI-A acknowledges funding from SECIHTI through scholarship 4070419. PEACCEL was supported through a research programme partially co-funded by the European Union (EU) and Région Réunion (FEDER).

## AI Statement

Generative artificial intelligence tools were used to support language editing, manuscript organisation, and refinement of explanatory text during the preparation of this work. These tools were not used to generate primary data, perform computational analyses, assign biological labels, select model configurations, calculate results, or draw scientific conclusions. All methodological decisions, analyses, interpretations, figures, tables, and final manuscript content were reviewed, verified, and approved by the authors, who take full responsibility for the accuracy and integrity of the work.

## References

Aguilera-Puga, M. D. C. and Plisson, F. (2024). Structure-aware machine learning strategies for antimicrobial peptide discovery. Scientific Reports, 14(1):11995.

Bernett, J., Blumenthal, D. B., Grimm, D. G., Haselbeck, F., Joeres, R., Kalinina, O. V., and List, M. (2024). Guiding questions to avoid data leakage in biological machine learning applications. Nature Methods, 21(8):1444–1453.

Bushuiev, A., Bushuiev, R., Sedlar, J., Pluskal, T., Damborsky, J., Mazurenko, S., and Sivic, J. (2024). Revealing data leakage in protein interaction benchmarks. arXiv (Cornell University).

Candido, S., Hayes, T., Derry, A., Rao, R., Lin, Z., Verkuil, R., Wu, B., Lee, J. S., Bruguera, E. S., Keval, J. A., Kopylov, M., Pak, J. E., Wu, W., Thomas, N., Mataraso, S., Hsu, A., Trotman-Grant, A. C., Fatras, K., dos Santos Costa, A., Badkundri, R., Akın, H., Oktay, D., Deaton, J., Montabana, E., Sitwala, H., Yu, Y., Wiggert, M., Carlin, D. A., Goering, A. W., Blazejewski, T., Sandora, M., Hla, M., Jia, T. Z., Kloker, L. H., Sofroniew, N. J., Uehara, M., Pannu, J., Bachas, S., Liu, D. S., Sercu, T., and Rives, A. (2026). Language modeling materializes a world model of protein biology. Preprint.

Chen, J. Y., Wang, J. F., Hu, Y., Li, X. H., Qian, Y. R., and Song, C. L. (2025). Evaluating the advancements in protein language models for encoding strategies in protein function prediction: a comprehensive review. Frontiers in Bioengineering and Biotechnology, 13:1506508.

Cheng, R., Liu, T., Liao, C., Wu, X., Zhu, L., and Zhang, S. (2026). Integrating Protein Language Models with Multimodal Embeddings to Accelerate Function Prediction of Uncharacterized Proteins. International Journal of Molecular Sciences, 27(9):3891.

Chhibbar, P. and Das, J. (2025). Machine learning approaches enable the discovery of therapeutics across domains. Molecular Therapy, 33(5):2269–2278.

Dickson, A. and Mofrad, M. R. K. (2024). Fine-tuning protein embeddings for functional similarity evaluation. Bioinformatics, 40(8).

Ekambaram, S. and Dokholyan, N. V. (2025). Peptide-based drug design using generative AI. Chemical Communications, 62(3):672–691.

Elnaggar, A., Essam, H., Salah-Eldin, W., Moustafa, W., Elkerdawy, M., Rochereau, C., and Rost, B. (2023). Ankh: Optimized protein language model unlocks general-purpose modelling. arXiv preprint arXiv:2301.06568.

Elnaggar, A., Heinzinger, M., Dallago, C., Rehawi, G., Wang, Y., Jones, L., Gibbs, T., Feher, T., Angerer, C., Steinegger, M., et al. (2021). Prottrans: toward understanding the language of life through self-supervised learning. IEEE transactions on pattern analysis and machine intelligence, 44(10):7112–7127.

Fernández-Díaz, R., Cossio-Pérez, R., Agoni, C., Lam, H. T., Lopez, V., and Shields, D. C. (2024). Autopeptideml: a study on how to build more trustworthy peptide bioactivity predictors. Bioinformatics, 40(9):btae555.

Graber, D., Stockinger, P., Meyer, F., Mishra, S., Horn, C., and Buller, R. (2025). Resolving data bias improves generalization in binding affinity prediction. Nature Machine Intelligence, 7(10):1713–1725.

Greener, J. G., Kandathil, S. M., Moffat, L., and Jones, D. T. (2022). A guide to machine learning for biologists. Nature reviews Molecular cell biology, 23(1):40–55.

Gujral, O., Bafna, M., Alm, E., and Berger, B. (2025). Sparse autoencoders uncover biologically interpretable features in protein language model representations. Proceedings of the National Academy of Sciences, 122(34):e2506316122.

Heil, B. J., Hoffman, M. M., Markowetz, F., Lee, S.-I., Greene, C. S., and Hicks, S. C. (2021). Reproducibility standards for machine learning in the life sciences. Nature Methods, 18(10):1132–1135.

Herrera-Rocha, F., Medina-Ortiz, D., Mauz, F., Pleiss, J., and Davari, M. D. (2025). Best Practices for Machine Learning-Assisted Protein Engineering. Journal of Chemical Information and Modeling, 65(23):12655–12667.

Hornback, A., Sathu, H., Kim, K., Wang, Y., Zhu, Y., Isgut, M., Avula, P., Khimani, A., and Wang, M. D. (2026). Large language models in healthcare and biomedical informatics: A comprehensive review. Innovation and Emerging Technologies, 13.

Hu, F., Hu, Y., Zhang, W., Huang, H., Pan, Y., and Yin, P. (2023). A multimodal protein representation framework for quantifying transferability across biochemical downstream tasks. Advanced Science, 10(22):e2301223.

Ibisanmi, T. A., Jiang, X., Willcox, M., and Kumar, N. (2026). Recent advances in computational antimicrobial peptide discovery through big data, modeling, and artificial intelligence and their interplay in ushering the next golden era of drug development. Frontiers in Bioinformatics, 6:1749404.

Jing, X., Dong, Q., Hong, D., and Lu, R. (2019). Amino acid encoding methods for protein sequences: a comprehensive review and assessment. IEEE/ACM transactions on computational biology and bioinformatics, 17(6):1918–1931.

Joeres, R., Blumenthal, D. B., and Kalinina, O. V. (2025). Data splitting to avoid information leakage with datasail. Nature Communications, 16(1):3337.

Kapoor, S. and Narayanan, A. (2023). Leakage and the reproducibility crisis in machine-learning-based science. Patterns, 4(9):100804.

Kucera, T., Oliver, C., Chen, D., and Borgwardt, K. (2023). Proteinshake: Building datasets and benchmarks for deep learning on protein structures. In Thirty-seventh Conference on Neural Information Processing Systems Datasets and Benchmarks Track.

Lin, B., Luo, X., Liu, Y., and Jin, X. (2024). A comprehensive review and comparison of existing computational methods for protein function prediction. Briefings in Bioinformatics, 25(4):bbae289.

Lin, Z., Akin, H., Rao, R., Hie, B., Zhu, Z., Lu, W., Smetanin, N., dos Santos Costa, A., Fazel-Zarandi, M., Sercu, T., Candido, S., et al. (2022). Language models of protein sequences at the scale of evolution enable accurate structure prediction. bioRxiv.

Liu, X., Zhang, J., Wang, X., Teng, M., Wang, G., and Zhou, X. (2025). Application of artificial intelligence large language models in drug target discovery. Frontiers in Pharmacology, 16:1597351.

Luque, A., Carrasco, A., Martín, A., and de Las Heras, A. (2019). The impact of class imbalance in classification performance metrics based on the binary confusion matrix. Pattern recognition, 91:216–231.

Martinez, K. M., Wilding, K., Llewellyn, T. R., Jacobsen, D. E., Montoya, M. M., Kubicek-Sutherland, J. Z., Batni, S., Manore, C., and Mukundan, H. (2025). Evaluating the factors influencing accuracy, interpretability, and reproducibility in the use of machine learning classifiers in biology to enable standardization. Scientific Reports, 15(1):16651.

Medina-Ortiz, D., Álvarez, D., and García-Vinuesa, J. (2026a). Sylphy: Protein sequence representation: Encoders, embeddings, and reductions. https://pypi.org/project/sylphy/. Python package, version 0.2.0, accessed 12 May 2026.

Medina-Ortiz, D., Álvarez-Saravia, D., and García-Vinuesa, J. (2026b). BioSieve: Redundancy reduction and leakage-aware dataset partitioning for biological machine learning. https://pypi.org/project/biosieve/. Python package, version 0.1.0, accessed 12 May 2026.

Medina-Ortiz, D., Escobedo, S., Murillo-Acevedo, N., Soto-García, N., Fernández-Villegas, D., Sandoval, D., and Daza, A. (2026c). Perspectives chapter: Data-centric strategies for machine learning-driven therapeutic peptide design – challenges and perspectives. In Ventura, S., Luna, J. M., and Martín-Castaño, A. R. M., editors, Data Quality Matters - Best Practices for Integrity and Assurance, chapter 23. IntechOpen, London.

Meng, C., Pei, Y., Bu, Y., Zou, Q., and Ju, Y. (2023). Machine learning-based antioxidant protein identification model: Progress and evaluation. Journal of Cellular Biochemistry, 124(11):1825–1834.

Mourad, R. (2024). Mistral-prot-v1-134m (mistral for protein). https://huggingface.co/RaphaelMourad/Mistral-Prot-v1-134M. Hugging Face Model Repository.

Pereira, J., Pantolini, L., Durairaj, J., and Schwede, T. (2025). Large-scale protein clustering in the age of deep learning. Current Opinion in Structural Biology, 94:103078.

Priyadarsinee, L., Kungurtsev, V., Kumar, V., Chatterjee, B., Sastry, G. N., and Murugan, N. A. (2026). Contemporary data-driven innovations in peptide-based therapeutic design. Briefings in Bioinformatics, 27(3).

Schütze, K., Heinzinger, M., Steinegger, M., and Rost, B. (2022). Nearest neighbor search on embeddings rapidly identifies distant protein relations. Frontiers in Bioinformatics, 2:1033775.

Shaw, R., Love, S. D., and McWhite, C. D. (2025). Evaluating pretrained protein language model embeddings as proxies for functional similarity. Journal of Molecular Evolution, 93(6):765–776.

Steinegger, M. and Söding, J. (2017). Mmseqs2 enables sensitive protein sequence searching for the analysis of massive data sets. Nature biotechnology, 35(11):1026–1028.

Teufel, F., Gíslason, M. H., Almagro Armenteros, J. J., Johansen, A. R., Winther, O., and Nielsen, H. (2023). Graphpart: homology partitioning for biological sequence analysis. NAR genomics and bioinformatics, 5(4):lqad088.

Tharwat, A. (2021). Classification assessment methods. Applied computing and informatics, 17(1):168–192.

Tran, C., Khadkikar, S., and Porollo, A. (2023). Survey of protein sequence embedding models. International Journal of Molecular Sciences, 24(4):3775.

Walsh, I., Fishman, D., Garcia-Gasulla, D., Titma, T., Pollastri, G., Harrow, J., Psomopoulos, F. E., and Tosatto, S. C. (2021). Dome: recommendations for supervised machine learning validation in biology. Nature methods, 18(10):1122–1127.

Walsh, I., Pollastri, G., and Tosatto, S. C. (2016). Correct machine learning on protein sequences: a peer-reviewing perspective. Briefings in bioinformatics, 17(5):831–840.

Wang, Y., Zhang, Y., Zhan, X., He, Y., Yang, Y., Cheng, L., and Alghazzawi, D. (2024). Machine learning for predicting protein properties: A comprehensive review. Neurocomputing, 597:128103.

Xiaoying, D., Kaijun, Y., Tianxiang, W., Pengyong, L., and Lin, G. (2026). A systematic benchmark for peptide property prediction. bioRxiv (Cold Spring Harbor Laboratory).

Xie, J., Li, Y., and Fu, T. (2025). DeepProtein: deep learning library and benchmark for protein sequence learning. Bioinformatics, 41(10).

Zha, D., Bhat, Z. P., Lai, K.-H., Yang, F., Jiang, Z., Zhong, S., and Hu, X. (2025). Data-centric artificial intelligence: A survey. ACM Computing Surveys, 57(5):1–42.

Zhang, J., Wang, R., Zhou, K., Xiao, T., Zhu, L., Min, Y., and Wang, Y. (2026). PePBenchmark: a standardized benchmark for peptide machine learning. arXiv (Cornell University).

Zhou, Q., Sun, L., Ou, Y., Liu, H., Gao, J., Dai, R., Li, Y., Cheng, G., Tao, Y., Pan, Y., and Huang, L. (2026). Artificial intelligence drives the identification and screening of novel antibiotics and antimicrobial peptides. Briefings in Bioinformatics, 27(2).

