## Supplementary Information for "Data-Centric Evaluation of Protein Function Prediction Pipelines"

#### Supplementary contents

**S1 Dataset construction and source curation** 2

#### S1 Dataset construction and source curation

This section describes the construction and curation of the antioxidant protein prediction dataset. It summarises how source-level dataset entries were identified, selected, retrieved, processed, and harmonised; how antioxidant and non-antioxidant labels were assigned; and how duplicate sequences and annotation conflicts were handled before downstream processing.

##### S1.1 Bibliographic search and source identification

Candidate sources were identified through a targeted bibliographic search focused on published datasets for antioxidant protein prediction. An initial exploratory search was conducted in September 2025 using the query *predict antioxidative activity of proteins.* This first screening was used to identify publications related to antioxidant protein prediction and to determine whether the associated sequence-level datasets were publicly accessible. At this stage, only a subset of the identified studies provided downloadable datasets, whereas other relevant articles were retained as contextual references but were not used as direct data sources because their datasets were not publicly available.

The search was subsequently expanded and updated between April and June 2026 using Google Scholar, web search engine queries, publisher pages, PubMed/PMC records, article supplementary materials, and linked code or data repositories. Search terms combined concepts related to the biological function, the prediction task, and the dataset format, including *antioxidant protein*, *AOP*, *antioxidant protein prediction*, *AOP prediction*, *non-antioxidant protein*, *protein antioxidant sequence classification*, *benchmark dataset*, *antioxidant classification models*, and *antioxidant proteins sequences*. Reference lists and dataset descriptions from relevant antioxidant protein prediction studies were also inspected to identify earlier benchmark datasets that had been reused, reformatted, or expanded in subsequent publications.

Table **S1** summarises the main search dimensions used during source identification. Priority was given to studies that provided sequence-level records in the article, supplementary material, GitHub repositories, or other associated repositories, and that reported labels compatible with a binary antioxidant/non-antioxidant prediction task.

**Supplementary Table S1**: Search strategy used to identify candidate sources for antioxidant protein dataset construction.

| **Search dimension** | **Example terms or sources** | **Purpose** |
|---|---|---|
| Initial exploratory search | *predict antioxidative activity of proteins* | Identify publications related to antioxidant protein prediction and assess whether the associated sequence-level datasets were publicly accessible. |
| Updated bibliographic search | Google Scholar, general web searches, publisher pages, PubMed/PMC records | Expand and update the initial search between April and June 2026 using bibliographic databases and publisher-indexed records. |
| Biological function | *antioxidant protein*, *AOP*, *antioxidant proteins sequences* | Identify studies focused on antioxidant protein function and sequence-level antioxidant annotations. |
| Prediction task | *antioxidant protein prediction*, *AOP prediction*, *non-antioxidant protein* | Recover studies framing antioxidant protein identification as a supervised binary classification problem. |
| Machine-learning and benchmark context | *protein antioxidant sequence classification*, *benchmark dataset*, *antioxidant classification models* | Prioritise studies that described machine-learning models, benchmark datasets, or sequence-based classification tasks. |
| Data availability screening | Article supplementary material, GitHub repositories, linked code repositories, and associated data repositories | Confirm whether sequence-level datasets and labels could be downloaded directly from sources provided by the authors. |
| Citation tracking | Reference lists and dataset descriptions from relevant antioxidant protein prediction studies | Identify earlier datasets reused, reformatted, recompiled, or expanded in subsequent publications. |

A candidate source-level dataset entry was retained for processing when sequence-level records were publicly accessible and the accompanying documentation allowed one or more of those records to be mapped to an antioxidant/non-antioxidant classification context. Retrieved entries could contain proteins, peptides, precursor sequences, or mixed sequence collections. Eligibility for inclusion in the analytical dataset was subsequently evaluated at the record level using predefined criteria for sequence availability, label interpretation, sequence quality, and length. Eligible records could be obtained from supplementary files, GitHub repositories, databases, the articles themselves, or other repositories indicated by the authors. Whenever possible, datasets were downloaded directly from the links provided in the corresponding publication, supplementary material, or associated repository.

Studies that discussed antioxidant prediction but did not provide accessible sequence-level records with interpretable labels were excluded from dataset processing, although they were retained for citation tracking and contextual review. Likewise, a retrieved source-level dataset entry could remain documented in the acquisition provenance even when none of its records ultimately satisfied the final eligibility criteria for inclusion in the protein-level analytical dataset.

When a publication or repository provided multiple files, dataset partitions, benchmark subsets, or supplementary tables, these files were treated as components of the same source-level dataset entry rather than as independent sources. All files associated with the same entry were retained during the initial compilation step to preserve the complete provenance of the retrieved data. This structure allowed within-entry consistency checks to be performed before cross-source integration, including cases in which the same sequence appeared in more than one file or partition associated with the same entry. The source-level citation, access date, file format, retrieval route, and file-specific origin were recorded to maintain traceability before subsequent record-level filtering, label standardisation, harmonisation, deduplication, consensus assignment, and redundancy-reduction procedures.

##### S1.2 Inclusion and exclusion criteria

Source-level dataset entries were retained for processing when sequence-level records were publicly accessible and the accompanying documentation allowed those records to be mapped to an antioxidant/non-antioxidant classification context. Retrieved entries could contain proteins, peptides, precursor sequences, or mixed sequence collections. Eligibility for inclusion in the analytical dataset was therefore evaluated primarily at the record level after source-specific parsing and label standardisation.

Records were retained for source-level standardisation and subsequent integration when they met all of the following criteria:

(i) an amino acid sequence was available from the article, a database, supplementary material, a GitHub repository, or another repository indicated by the authors;

(ii) the record could be mapped to an antioxidant or non-antioxidant classification context using the original source documentation;

(iii) the antioxidant or non-antioxidant label was reported explicitly, or the source provided a clear definition from which the binary label could be interpreted; and

(iv) the record could be converted into a structured sequence–label representation while preserving its source-level provenance.

Individual records were excluded during source-specific processing when the sequence was missing or malformed, the label could not be interpreted unambiguously, the record contained only aggregate information, or the available information could not be converted into a structured sequence–label table. Source-level dataset entries were excluded from processing only when no accessible records satisfied the minimum sequence and label requirements.

Final eligibility for the protein-level analytical dataset was determined after cross-source integration and consensus assignment. At that stage, sequences containing non-canonical amino acids or falling outside the accepted length range of 70–1024 residues were removed. Consequently, a source-level dataset entry could remain documented in the acquisition and provenance records even when some or all of its retrieved sequences were excluded from the final analytical dataset.

##### S1.3 Source-level dataset entries retained for processing

This subsection summarises the source-level dataset entries retained during acquisition and source-specific processing. Table **S2** reports, for each entry, the retrieved files, access route and date, original sequence type, number of retrieved records, number of unique sequences obtained after initial record-level processing, and the distribution of antioxidant and non-antioxidant labels interpreted from the corresponding source.

The sequence type reported in the table describes the biological scope of the retrieved resource or collection and does not imply that every record satisfied the criteria subsequently used to define the protein-level analytical dataset. The retained source-level dataset entries included protein, peptide, and mixed peptide/protein collections. In particular, AMPDB v1 contained both peptide and protein records associated with multiple biological activities, whereas the AOPxSVM entry corresponded specifically to antioxidant-peptide training and evaluation collections. The positive and negative record counts preserve the labels provided by each original source and therefore refer to source-level records rather than exclusively to proteins retained in the final analytical dataset.

Retrieved-record and unique-sequence counts are specific to each source-level dataset entry and are not mutually exclusive across entries, because identical amino acid sequences may be reported by more than one source. Consequently, these values should not be summed to obtain the number of globally unique sequences. After cross-source integration, consensus assignment, exclusion of conflicting annotations, canonical-residue filtering, and application of the accepted length range of 70–1024 residues, the final curated consensus protein dataset contained 4,193 globally unique sequences, including 1,010 antioxidant and 3,183 non-antioxidant proteins.

The *Final set?* column indicates whether at least one sequence reported by the corresponding source-level dataset entry remained represented in the final protein-level analytical dataset. This binary indicator does not represent an additive or mutually exclusive per-source contribution. AOPxSVM [Li et al., 2025] was retained in the acquisition provenance because its publicly available files could be retrieved, its sequences could be processed, and its antioxidant and non-antioxidant labels could be interpreted. However, the entry consisted of antioxidant-peptide collections, and none of its 3,144 unique processed sequences remained after application of the length eligibility criteria used to define the final protein-level dataset. AOPxSVM is therefore documented as a retrieved and processed source-level dataset entry but is reported as *No* in the *Final set?* column.

Ho/Thanh-Lam et al. [Ho Thanh Lam et al., 2020] appears in two rows because two source-level dataset entries were retrieved and processed separately. These entries differed in access route, retrieval date, supplementary-data provenance, and reported sequence origin. One entry was retrieved from the PMC-linked supplementary material and included data described as derived from the independent dataset of Butt et al. [2019], whereas the second entry was retrieved from the publisher's supplementary material and was described as derived from Zhang et al. [2016] and Fernández-Blanco et al. [2013].

**Supplementary Table S2**: Source-level dataset entries retained during acquisition and source-specific processing.

| Source | Retrieved file(s) | Route | Sequence type | Access | Retrieved records | Unique after processing | Positive records | Negative records | Final set? |
|---|---|---|---|---|---|---|---|---|---|
| Ahmad et al. [2022] | `Training_dataset.txt`<br>`Indpndet_dataset.txt` | GitHub | Protein | Apr-26 | 665 | 665 | 173 | 492 | Yes |
| AMPDB v1 [Mondal et al., 2023] | `Antioxidantdataset.tsv` | Database | Peptide/ protein | Aug-25 | 356 | 344 | 344 | 0 | Yes |
| ANOX [Sun et al., 2021] | `anti_protein_positive_negative.txt` | GitHub | Protein | Sep-25 | 1,805 | 1,805 | 253 | 1,552 | Yes |
| AOD [Feng et al., 2017] | 36 timestamped files<br>`20260408*.xls` | Database | Protein | Apr-26 | 710 | 651 | 651 | 0 | Yes |
| AOPxSVM [Li et al., 2025] | `AOPP.train.fasta`<br>`AOPP.test.fasta`<br>`AOPP.test.2023.fasta` | GitHub | Peptide | Apr-26 | 3,172 | 3,144 | 1,576 | 1,568 | **No** |
| Butt et al. [2019] | `1-s2.0-S0022519319301602-mmc1.xlsx` | Publisher SI | Protein | Sep-25 | 1,945 | 1,945 | 370 | 1,575 | Yes |
| Feng et al. [2013] | `567529.f1.pdf` | PMC SI | Protein | Apr-26 | 1,841 | 1,840 | 274 | 1,566 | Yes |
| Ho Thanh Lam et al. [2020] PMC entry | `data.xlsx` | PMC SI | Protein | Sep-25 | 2,366 | 2,322 | 399 | 1,923 | Yes |
| PredAoDP [Ahmed et al., 2022] | `anti.txt`<br>`nonanti.txt`<br>`Antioxident666.txt`<br>`independentdataset.txt`<br>`Indpendentdataset.txt` | GitHub | Protein | Sep-25 | 3,007 | 2,378 | 365 | 2,013 | Yes |
| Ho Thanh Lam et al. [2020] Publisher entry | `data.xlsx` | Publisher SI | Protein | Apr-26 | 466 | 466 | 74 | 392 | Yes |
| Zhai et al. [2020] | `anti.txt`<br>`nonanti.txt` | GitHub | Protein | Sep-25 | 1,805 | 1,805 | 253 | 1,552 | Yes |
| Zhang et al. [2016] | `pone.0163274.s001` | Publisher SI | Protein | Sep-25 | 666 | 666 | 174 | 492 | Yes |

##### S1.4 Source-level parsing, metadata standardisation, and label assignment

After the source-level dataset entries were retained, each entry was processed with a dedicated parsing notebook adapted to the format and organisation of the retrieved files. The input files varied across sources and included plain-text files, FASTA-like records, tabular files, spreadsheets, and supplementary datasets distributed as training, testing, independent, positive, or negative subsets. When an entry was distributed across multiple files or dataset partitions, all files associated with that entry were parsed together and treated as components of the same source-level dataset entry. The overall source-level parsing workflow is summarised in Figure **S1**.

For each source-level dataset entry, amino acid sequence records were extracted and converted into a standardised sequence-level table. Labels were assigned according to the information provided by the original source, including file names, sheet names, sequence identifiers, class columns, dataset partitions, or dataset descriptions. Positive examples corresponded to records reported by the source as antioxidant or as members of the positive antioxidant class. Negative examples corresponded to records reported as non-antioxidant, inactive, or as members of the negative class defined by the original authors. The standardised output of each parser consisted of at least two fields, `sequence` and `label`. This source-level standardisation was performed before cross-source harmonisation and final protein-level sequence filtering; therefore, the labels at this stage represented the interpretation of each individual source rather than a final consensus label.

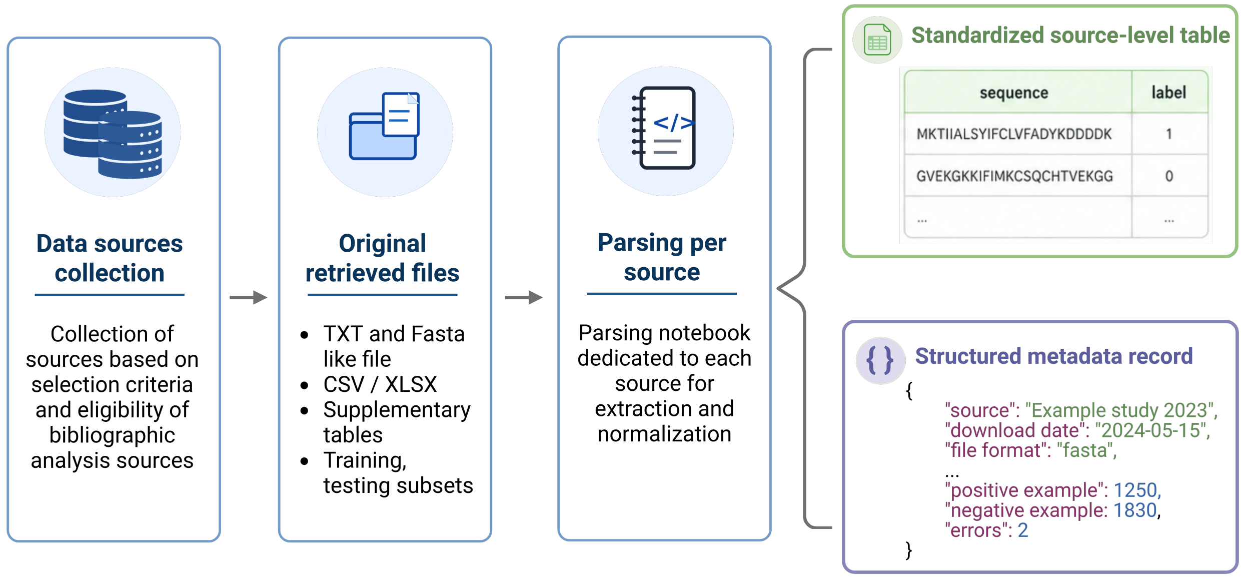

**Supplementary Figure S1**: **Source-level parsing and metadata standardisation workflow.** Retained source-level dataset entries were processed using dedicated parsing notebooks adapted to the organisation and format of the original files. Each parser produced a standardised source-level sequence-label table and a structured metadata record documenting provenance, file characteristics, label counts, and parsing outcomes. These outputs were used as inputs for the subsequent cross-source harmonisation step.

The rules used for source-level label assignment are summarised in Table **S3**. No label was reassigned based on sequence similarity, external functional assumptions, or downstream model behaviour at this stage. Records were retained only when their source-level label could be interpreted unambiguously as antioxidant or non-antioxidant.

**Supplementary Table S3**: Rules used for source-level positive and negative label assignment before cross-source harmonisation.

| **Source information** | **Assignment rule** | **standardised label** |
|---|---|---|
| Explicit class column | Labels were mapped directly when the source provided antioxidant or non-antioxidant annotations. | Positive or negative according to the reported class. |
| Separate positive and negative files | Labels were inferred from file names, dataset partitions, or accompanying documentation. | Positive for antioxidant files and negative for non-antioxidant files. |
| Training, testing, or independent subsets | Original class definitions were preserved across all subsets belonging to the same source. | Positive or negative according to the source definition. |
| Positive-only datasets | Sequences were retained when explicitly reported as antioxidant proteins. | Positive only. |
| Ambiguous or missing labels | Records were not assigned a binary label when the class could not be interpreted unambiguously. | Excluded from standardised source-level tables. |

During parsing, malformed entries, missing sequence records, empty labels, and sequence strings that could not be interpreted as valid amino acid sequence records were identified and removed when detected. Source-level dataset entries containing only antioxidant-labelled records were retained as positive-only evidence entries when the sequences and labels were explicitly available. These entries contributed positive source-level evidence but did not provide negative examples. Conversely, entries containing both antioxidant- and non-antioxidant-labelled records contributed both positive and negative evidence according to the original class definitions.

In parallel, a metadata file was generated for each processed source. These metadata records documented both provenance and processing information, including the source name, dataset file names, source type, static or dynamic status, licence information when available, publication year, last update date, download date, file format, protein representation format, task, repository or publication link, and the criteria used by the original source to define positive and negative examples. Processing-level fields were also recorded, including the processing date, number of retrieved records, number of collected sequences, number of unique sequences after initial processing, number of positive and negative examples, and number of sequences with parsing or formatting errors.

The output of this step was therefore a collection of standardised source-level datasets accompanied by structured metadata. These files provided the input for the subsequent cross-source integration step, in which labels from multiple sources were aligned, source-specific annotations were compared, duplicate sequences were evaluated, and annotation conflicts were detected.

##### S1.5 Cross-source integration, consensus assignment, and final sequence filtering

After source-level parsing and label assignment, the standardised datasets were integrated into a sequence-centred evidence table. For each retained source-level dataset entry, the corresponding `processed_data.csv` file was loaded, annotated with its source-entry identifier, and concatenated into a single table containing the fields `sequence`, `label`, and `source`. This table was then reshaped into a pivot format, in which each row represented a unique amino acid sequence and each source-level dataset entry was represented as a separate column containing the label assigned by that entry.

Missing source-entry–sequence combinations were treated as absence of evidence and were not interpreted as either antioxidant or non-antioxidant labels. For each sequence, the number of source-level dataset entries assigning antioxidant and non-antioxidant labels was computed and normalised by the number of entries reporting that sequence. These proportions were used to assign consensus labels. Sequences for which all reporting source-level dataset entries assigned the antioxidant label were retained as consensus-positive examples, whereas sequences for which all reporting entries assigned the non-antioxidant label were retained as consensus-negative examples. Thus, consensus labels were operationally defined by complete agreement among all source-level dataset entries reporting a given sequence.

Sequences with mixed evidence, *i.e.*, sequences assigned as antioxidant by at least one source-level dataset entry and as non-antioxidant by at least one other entry, were not forced into either class. Instead, they were separated into a conflict file for traceability and excluded from the curated consensus binary dataset. The merging step therefore generated three main outputs: a consensus-labelled dataset, a file containing sequences with conflicting annotations, and a metadata record summarising the number of integrated sequences, conflicting cases, uniquely annotated sequences, consensus-positive and consensus-negative examples, and the observed sequence-length range before final filtering. Source-entry support was recorded as the number of source-level dataset entries contributing concordant evidence for the consensus label assigned to each sequence.

Final record-level quality control was applied in a separate preprocessing step after cross-source integration and consensus assignment. This step defined the protein-level analytical dataset used in downstream experiments. Sequence length was computed when needed, canonical amino acid composition was evaluated using the predefined standard residue vocabulary, and length constraints were applied using the established minimum and maximum sequence-length thresholds. The minimum length threshold was set to 70 residues to exclude short peptide-like records from the protein classification task, whereas the maximum threshold was set to 1024 residues according to the sequence-length restrictions of the selected protein language models.

Both the consensus-labelled dataset and the conflict file were evaluated using the same quality-control rules. Sequences were retained in the corresponding filtered output only when they contained canonical amino acid residues and fell within the accepted length range. The preprocessing step exported the final curated consensus protein dataset, a filtered conflict file, and a metadata record summarising the number of sequences before and after filtering, the number of canonical and non-canonical entries, the number of sequences inside and outside the accepted length range, and the final distribution of positive and negative examples. Source-level dataset entries and excluded records remained documented in the provenance and processing metadata even when they did not contribute sequences to the final analytical dataset.

The rules applied during cross-source integration and final filtering are summarised in Table **S4**.

##### S1.6 Final dataset composition after harmonisation and filtering

After source-level parsing, cross-source integration, consensus-based label assignment, conflict detection, and final sequence-level quality control, the curated antioxidant protein dataset was summarised to document

**Supplementary Table S4**: Rules applied during cross-source integration, consensus label assignment, conflict detection, and final record-level sequence filtering.

| Processing case | Rule applied | Output or interpretation |
|---|---|---|
| Sequence absent from a source-level dataset entry | The source-level dataset entry did not report the sequence. | Treated as absence of evidence, not as a negative label. |
| Sequence reported by one or more source-level dataset entries with only antioxidant labels | All valid source-level labels reported for the sequence were positive. | Assigned to the consensus-positive class. |
| Sequence reported by one or more source-level dataset entries with only non-antioxidant labels | All valid source-level labels reported for the sequence were negative. | Assigned to the consensus-negative class. |
| Sequence reported with both antioxidant and non-antioxidant labels across source-level dataset entries | At least one source-level dataset entry assigned a positive label and at least one other entry assigned a negative label. | Flagged as a conflicting annotation and exported separately from the curated consensus binary dataset. |
| Repeated sequence across different source-level dataset entries | Identical sequences were merged into a single row in the evidence matrix while preserving each source-level label. | Used for consensus assignment, conflict detection, and source-entry support calculation. |
| Sequence containing non-canonical residues | The sequence failed the canonical amino acid vocabulary check. | Removed during final record-level quality control. |
| Sequence outside the accepted length range | The sequence contained fewer than 70 residues or more than 1024 residues. | Excluded from the protein-level analytical dataset during final record-level quality control. |
| Sequence passing both canonical-residue and length criteria | The sequence contained only canonical amino acids and fell within the accepted range of 70–1024 residues. | Retained in the corresponding filtered output; consensus-labelled sequences formed the final curated consensus protein dataset. |

the effect of each processing step. Table **S5** reports the number of records or unique sequences retained at the main stages of dataset construction, including source-level inputs, cross-source integration, ambiguous annotations, filtering outcomes, and the final curated consensus protein dataset used for downstream analyses. These counts provide an audit trail of the curation process and complement the dataset-level analyses reported in the main Results section. Counts reported for the canonical-residue and length filters correspond to independent evaluations of each quality-control criterion. Consequently, individual sequences may contribute to both filtering categories, and these counts should not be interpreted as mutually exclusive.

**Supplementary Table S5**: Final antioxidant protein dataset summary after harmonisation and filtering. Counts summarise the main stages of source integration, annotation conflict detection, and sequence-level quality control.

| Curation stage | Total | Positive | Negative | Conflict |
|---|---|---|---|---|
| Retrieved source-level records | 18,804 | – | – | - |
| Unique sequences after source-level processing | 18,031 | 4,906 | 13,125 | 0 |
| Unique sequences after cross-source integration | 7,870 | 2,649 | 5,178 | 43 |
| Consensus-labelled sequences before final filtering | 7,827 | 2,649 | 5,178 | 0 |
| Removed due to non-canonical residues | 53 | 27 | 26 | 0 |
| Removed due to length filtering | 3,609 | 1,613 | 1,993 | 3 |
| Final curated consensus protein dataset | 4,193 | 1,010 | 3,183 | 0 |

#### S1.7 Source-support structure of the final consensus dataset

In addition to the final consensus label, source-entry support was computed as the number of source-level dataset entries contributing concordant labels for each retained sequence. This count was used to summarise the documentary evidence structure of the final dataset and to define source-support-based subsets for sensitivity analysis, as described in Supplementary Section S7.

Source-entry support should not be interpreted as the number of independent experiments or independent biological validations. Source-level dataset entries may reuse benchmark records, redistribute earlier collections, or share upstream dataset lineages. Accordingly, the measure quantifies concordant reporting across retrieved entries and does not provide a direct estimate of label reliability or experimental replication.

**Supplementary Table S6**:
Source-support distribution in the final curated consensus protein dataset.

| Supporting source-level dataset entries | Total sequences | Antioxidant proteins | Non-antioxidant proteins | Antioxidant (%) | Non-antioxidant (%) |
|---|---|---|---|---|---|
| 1 | 1,895 | 573 | 1,322 | 30.2 | 69.8 |
| 2 | 91 | 90 | 1 | 98.9 | 1.1 |
| 3 | 90 | 39 | 51 | 43.3 | 56.7 |
| 4 | 34 | 32 | 2 | 94.1 | 5.9 |
| 5 | 1,807 | 5 | 1,802 | 0.3 | 99.7 |
| $\geq 6$ | 276 | 271 | 5 | 98.2 | 1.8 |
| Overall | 4,193 | 1,010 | 3,183 | 24.1 | 75.9 |

### S2 Protein sequence representations

This section describes the numerical representations used to encode the curated antioxidant protein sequences before redundancy reduction, dataset splitting, and model training. The study included one sequence-derived baseline representation, based on one-hot encoding, and six pretrained protein language model representations.

#### S2.1 Numerical representation extraction protocol

Numerical sequence representations were generated from the harmonised antioxidant protein sequences using the Sylphy framework [Medina-Ortiz et al., 2026a]. The same curated input sequence set was used for all representation types to ensure that downstream comparisons reflected differences in the numerical encoding strategy rather than differences in the encoded sequences. Two representation-generation modes were used: pretrained protein language model embeddings and one-hot sequence encoding.

For protein language models, embeddings were extracted using pretrained models without task-specific fine-tuning. Residue-level representations were obtained from the final hidden layer, and fixed-length sequence-level vectors were generated by applying mean pooling across residue embeddings. Embedding extraction was performed with `cuda` as the computation device, `fp32` precision, a batch size of 16, and a maximum sequence length of 1024 residues.

For protein language models, residue-level embeddings from the final hidden layer were denoted as

$$H_i = (h_{i1}, h_{i2}, \dots, h_{iL_i}), \quad h_{il} \in \mathbb{R}^d,$$

where $L_i$ is the sequence length of protein $i$, $d$ is the embedding dimension of the corresponding protein language model, and $h_{il}$ is the residue-level embedding for residue $l$. The sequence-level embedding was obtained by mean pooling across residues:

$$z_i = \frac{1}{L_i} \sum_{l=1}^{L_i} h_{il}.$$

For the one-hot baseline, sequences were encoded directly from the amino acid alphabet using Sylphy's one-hot encoding mode. Unlike the pretrained model representations, this representation does not involve a learned embedding layer or pooling operation. Each residue was mapped to a binary vector over the 20 canonical amino acids. Sequences shorter than $L_{\max} = 1024$ were zero-padded, producing fixed-size sequence-derived vectors suitable for downstream classical machine-learning models.

All numerical representations were exported in CSV format. For each representation, companion files were retained to preserve the correspondence between each numerical vector, the original protein sequence, and the associated label used in downstream analyses. The representation extraction settings are summarised in Table **S7**.

**Supplementary Table S7**: Settings used to generate numerical representations for antioxidant protein sequences. Both pretrained protein language model embeddings and one-hot sequence encodings were generated using Sylphy.

| **Parameter** | **Protein language model embeddings** | **One-hot encoding** |
|---|---|---|
| Framework | Sylphy | Sylphy |
| Sylphy version | v0.2.0 | v0.2.0 |
| Representation method | `sylphy_embedding` | `sylphy_one_hot` |
| Model or encoder | Pretrained protein language model | `one_hot` encoder |
| Model state | Pretrained, without task-specific fine-tuning | Not applicable |
| Input sequences | harmonised antioxidant protein sequences after dataset curation and filtering | harmonised antioxidant protein sequences after dataset curation and filtering |
| Representation layer | final hidden layer | Not applicable |
| Pooling strategy | Mean pooling across residue-level embeddings | Not applicable |
| Device | `cuda` | Not applicable; encoder-based representation |
| Precision | `fp32` | Not applicable |
| Batch size | 16 | Not applicable |
| Maximum sequence length | 1024 residues | 1024 residues |
| Output format | CSV | CSV |

##### S2.2 Representation inventory

Table **S8** summarises the sequence representations used in the core experiments. For protein language models, the table reports the model identifier used for extraction, the model family, the numerical dimensionality, and the pooling strategy used to obtain one fixed-length vector per sequence. The one-hot representation was generated directly from the amino acid sequence alphabet and was included as a non-pretrained baseline.

**Supplementary Table S8**: Protein sequence representations used to encode antioxidant protein sequences. The table reports the representation name, model identifier or origin, model family, numerical dimensionality, pooling strategy, and reference used for each representation.

| **Representation** | **Model identifier or origin** | **Architecture / family** | **Dimension** | **Pooling** | **Reference** |
|---|---|---|---|---|---|
| One-hot | Sequence-derived encoding | Amino acid alphabet encoding | 20480 ($1024 \times 20$) | Not applicable | [Jing et al., 2019] |
| Ankh2-ext1 | `ElnaggarLab/ankh2-ext1` | Ankh protein language model | 1536 | Mean pooling | [Elnaggar et al., 2023] |
| ESM2-8M | `facebook/esm2_t6_8M_UR50D` | ESM2 transformer | 320 | Mean pooling | [Lin et al., 2022] |
| ESMC-300M | `esmc_300m` | ESM Cambrian model | 960 | Mean pooling | [Candido et al., 2026] |
| Mistral-Prot | `RaphaelMourad/Mistral-Prot-v1-134M` | Causal protein language model | 768 | Mean pooling | [Mourad, 2024] |
| ProtBERT | `Rostlab/prot_bert` | ProtTrans BERT | 1024 | Mean pooling | [Elnaggar et al., 2021] |
| ProtT5-XL | `Rostlab/prot_t5_xl_uniref50` | ProtTrans T5 encoder | 1024 | Mean pooling | [Elnaggar et al., 2021] |

#### S3 Redundancy reduction procedures

To reduce the probability that identical, homologous, or highly similar sequences were distributed across training, validation, and test sets, redundancy control was applied before dataset splitting and model training. This step was designed to reduce the risk of *data leakage* and to provide a stricter assessment of model generalisation [Bernett et al., 2024].

Two complementary notions of sequence similarity were considered. The first was sequence homology, estimated from amino acid sequence identity, which captures redundancy from a classical biological sequence-comparison perspective [Teufel et al., 2023]. The second was similarity in numerical representation spaces, where two sequences were considered close when their encoded vectors showed high similarity or low distance

[Schütze et al., 2022]. Because this second criterion depends on the geometry induced by each representation, representation-distance-based reduction was interpreted separately for each encoding space.

Both reduction strategies were implemented using `BioSieve` v0.1.0 [Medina-Ortiz et al., 2026b]. Homology-based reduction used the `MMseqs2` backend available through `BioSieve`, with `MMseqs2` version 18.8cc5c [Steinegger and Söding, 2017]. Representation-distance-based reduction used the numerical representations generated for the curated antioxidant protein dataset. The general settings used for both reduction families are summarised in Table **S9**.

**Supplementary Table S9**: Settings used for homology-based and representation-distance-based redundancy reduction.

| **Parameter** | **Homology-based reduction** | **Representation-distance-based reduction** |
|---|---|---|
| Workflow | `BioSieve` v0.1.0 | `BioSieve` v0.1.0 |
| Reduction strategy | `mmseqs2` | `embedding_cosine` for protein language model embeddings; `descriptor_euclidean` for one-hot when applicable |
| External backend | `MMseqs2` v18.8cc5c | Not applicable for embedding-space reduction |
| Reduction criterion | Amino acid sequence identity | Pairwise similarity or distance in numerical representation space |
| Metric | Minimum sequence identity with minimum coverage | Cosine similarity for protein language model embeddings; Euclidean distance for one-hot descriptor vectors when applicable |
| Threshold definition | Fixed minimum sequence identity thresholds | Representation-specific thresholds derived from empirical percentile tables |
| Thresholds evaluated | 0.9, 0.7, 0.5, and 0.3 minimum sequence identity | Percentile-based thresholds read from the corresponding representation-specific similarity or distance percentile table |
| Coverage | 0.8 minimum coverage | Not applicable |
| Representative selection | One representative sequence retained per redundancy group using the default `BioSieve` selection rule | One representative sequence retained per redundancy group using the default `BioSieve` selection rule |
| Recorded outputs | Reduced dataset, sequence-to-representative mapping, reduction report, retained/removed counts, and label distribution | Reduced dataset, sequence-to-representative mapping, reduction report, retained/removed counts, and label distribution |

#### S3.1 Homology-based redundancy reduction

Homology-based redundancy reduction was performed to remove sequences sharing high amino acid sequence identity. This reduction was carried out with the `mmseqs2` strategy implemented in `BioSieve` v0.1.0 [Medina-Ortiz et al., 2026b], using `MMseqs2` v18.8cc5c. Four minimum sequence identity thresholds were evaluated: 0.9, 0.7, 0.5, and 0.3. A minimum coverage of 0.8 was used in all cases, with `cov_mode = 0`, `cluster_mode = 0`, and 16 threads. Each threshold generated an independently reduced dataset, allowing the effect of increasingly stringent homology control to be evaluated. Lower identity thresholds correspond to stricter redundancy reduction because they collapse sequences at lower levels of sequence identity.

Homology-based reduction was performed directly on the protein sequence column, rather than on numerical representation distances. At each identity threshold, sequences were clustered according to the MMseqs2 sequence-similarity criterion, and a single representative sequence was retained from each redundancy group. Representative selection was handled by `MMseqs2 easy-cluster` through the `BioSieve` workflow.

The outputs of each homology-reduction run included the reduced dataset, a mapping file linking removed sequences to retained representatives, a reduction report, and a summary table. The number of retained and removed sequences was recorded for each threshold. Label-specific retention was evaluated separately during the reduction summary and label-aware analysis step, using the labelled reduced datasets generated by the workflow. The homology-reduction results are summarised in Table **S10**.

#### S3.2 Representation-distance-based redundancy reduction

Representation-distance-based redundancy reduction was performed to evaluate redundancy according to the geometry induced by each numerical representation. In this approach, the reduction criterion was not amino acid identity, but proximity between encoded sequence vectors. For protein language model embeddings,

**Supplementary Table S10**: Dataset size after homology-based redundancy reduction with `MMseqs2`.

| Identity threshold | Input sequences | Retained sequences | Removed sequences | Retained (%) | Removed (%) |
|---|---|---|---|---|---|
| 0.9 | 4193 | 3801 | 392 | 90.65 | 9.35 |
| 0.7 | 4193 | 3510 | 683 | 83.71 | 16.29 |
| 0.5 | 4193 | 3252 | 941 | 77.56 | 22.44 |
| 0.3 | 4193 | 2948 | 1245 | 70.31 | 29.69 |

redundancy was evaluated using cosine similarity. For the one-hot representation, descriptor-space reduction was evaluated using Euclidean distance between one-hot-derived vectors.

Cosine similarity between two sequence embeddings $z_i$ and $z_j$ was computed as

$$\cos(z_i, z_j) = \frac{z_i z_j}{\|z_i\|\|z_j\|}.$$

For one-hot descriptor vectors, Euclidean distance was computed as

$$d(x_i, x_j) = \sqrt{\sum_{k=1}^{p}(x_{ik} - x_{jk})^2},$$

where $p$ is the dimensionality of the one-hot-derived descriptor vector.

For each representation $r$, percentile thresholds were derived from the empirical distribution of pairwise similarities or distances:

$$\tau_{r,p} = Q_p(\mathcal{D}_r),$$

where $Q_p$ denotes the $p$-th percentile and $\mathcal{D}_r$ is the set of pairwise similarity or distance values computed in representation space $r$. Because $\mathcal{D}_r$ is representation-specific, percentile thresholds were interpreted within each representation space rather than as directly comparable absolute cutoffs across representations.

For embedding-based reductions, thresholds were obtained from the empirical distribution of pairwise cosine similarities computed independently for each representation. Specifically, reduction thresholds were read from the representation-specific percentile table generated during representation-space analysis. The evaluated percentiles were p30, p40, p50, p60, p70, p80, p90, p95, p97, p98, p99, p99.5, and p99.9. Because each percentile corresponds to a different absolute similarity value depending on the embedding model, the same percentile should not be interpreted as the same redundancy threshold across representations. Instead, each percentile defines a reduction level relative to the geometry of the corresponding representation space.

Before applying distance-based reduction, the geometry of each embedding space was characterised using cosine-similarity distributions. As shown in Supplementary Figure **S2**, the evaluated representations differed markedly in the distribution of pairwise similarities. Ankh2-ext1 and ProtT5-XL showed broader distributions shifted towards lower similarity values, whereas ESM2-8M, ESMC-300M, ProtBERT, and especially Mistral-Prot concentrated a larger fraction of sequence pairs at high cosine similarity. These differences indicate that each embedding model imposes a distinct organisation on the same sequence dataset. Consequently, percentile-based reduction was interpreted within each representation space rather than as a universal similarity cutoff shared across embeddings.

In embedding-based reductions, the representation used to compute redundancy was distinguished from the representation later used as input to the classifier. This distinction allowed the effect of the reduction space to be evaluated separately from the effect of the predictive representation used during model training. For each representation and percentile threshold, sequences satisfying the redundancy criterion were grouped, and one representative sequence was retained per group using the default `BioSieve` selection rule.

For each reduced dataset, the number of retained and removed sequences was recorded together with the corresponding antioxidant and non-antioxidant label distribution. These summaries were used to assess both the intensity of redundancy reduction and its effect on class balance. The outcomes of representation-distance-based reduction are summarised in Table **S11**. Because the one-hot representation did not lead to sequence removal at any evaluated percentile, its class distribution remained identical to that of the input dataset and is therefore summarised once rather than repeated across thresholds.

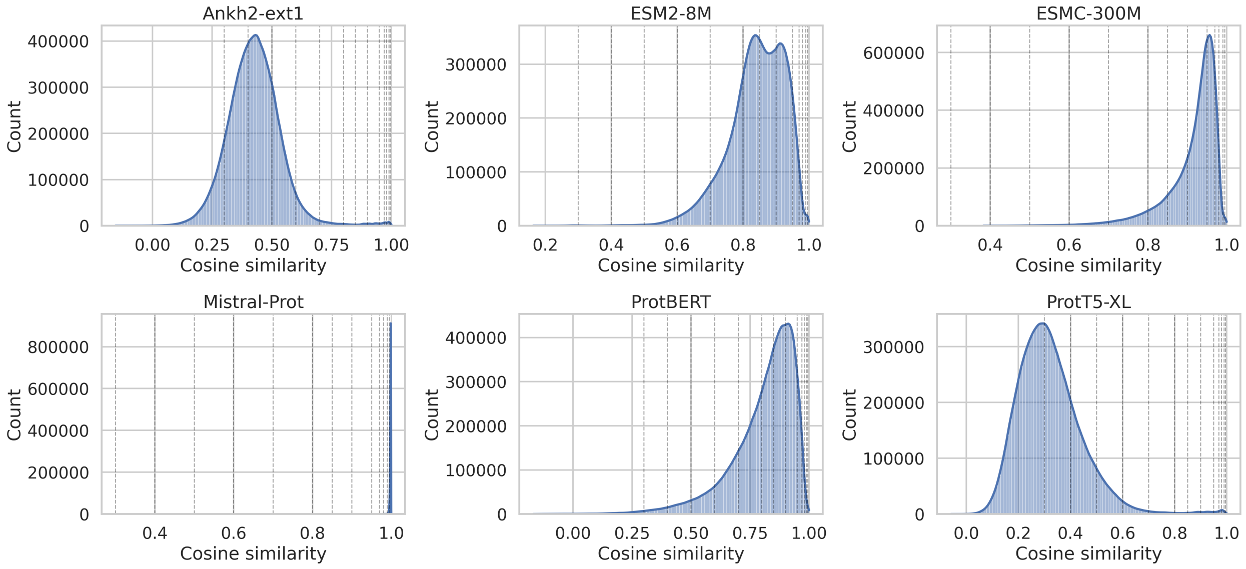

**Supplementary Figure S2**: Cosine-similarity distributions across protein language model embeddings used for representation-distance-based redundancy reduction. Vertical dashed lines indicate the evaluated percentile thresholds. The distributions show that different embeddings induce distinct similarity geometries over the same antioxidant protein dataset; therefore, percentile-based thresholds were interpreted within each representation space rather than as directly comparable absolute similarity cutoffs.

**Supplementary Table S11**: Representation-distance-based redundancy reduction summary by percentile. For protein language model embeddings, thresholds correspond to representation-specific cosine similarity values derived from empirical percentile distributions. For one-hot, Euclidean-distance thresholds were evaluated across the same percentile range, but no sequences were removed. Antioxidant and non-antioxidant counts correspond to retained class counts after reduction.

| **Reduction space** | **Percentile** | **Threshold** | **Retained** | **Removed** | **Antioxidant** | **Non-antioxidant** |
|---|---|---|---|---|---|---|
| Ankh2-ext1 | p30 | 0.3715 | 24 | 4169 | 2 | 22 |
| Ankh2-ext1 | p40 | 0.3985 | 30 | 4163 | 2 | 28 |
| Ankh2-ext1 | p50 | 0.4237 | 59 | 4134 | 4 | 55 |
| Ankh2-ext1 | p60 | 0.4484 | 95 | 4098 | 5 | 90 |
| Ankh2-ext1 | p70 | 0.4750 | 151 | 4042 | 10 | 141 |
| Ankh2-ext1 | p80 | 0.5060 | 261 | 3932 | 14 | 247 |
| Ankh2-ext1 | p90 | 0.5505 | 453 | 3740 | 20 | 433 |
| Ankh2-ext1 | p95 | 0.5930 | 708 | 3485 | 30 | 678 |
| Ankh2-ext1 | p97 | 0.6283 | 971 | 3222 | 44 | 927 |
| Ankh2-ext1 | p98 | 0.6615 | 1241 | 2952 | 58 | 1183 |
| Ankh2-ext1 | p99 | 0.7553 | 1935 | 2258 | 87 | 1848 |
| Ankh2-ext1 | p99.5 | 0.9066 | 2926 | 1267 | 191 | 2735 |
| Ankh2-ext1 | p99.9 | 0.9872 | 3652 | 541 | 503 | 3149 |
| ESM2-8M | p30 | 0.8071 | 114 | 4079 | 6 | 108 |
| ESM2-8M | p40 | 0.8294 | 141 | 4052 | 7 | 134 |
| ESM2-8M | p50 | 0.8502 | 181 | 4012 | 12 | 169 |
| ESM2-8M | p60 | 0.8724 | 246 | 3947 | 17 | 229 |
| ESM2-8M | p70 | 0.8953 | 338 | 3855 | 22 | 316 |
| ESM2-8M | p80 | 0.9173 | 483 | 3710 | 33 | 450 |
| ESM2-8M | p90 | 0.9403 | 726 | 3467 | 56 | 670 |
| ESM2-8M | p95 | 0.9547 | 1041 | 3152 | 83 | 958 |
| ESM2-8M | p97 | 0.9624 | 1294 | 2899 | 120 | 1174 |
| ESM2-8M | p98 | 0.9675 | 1540 | 2653 | 148 | 1392 |
| ESM2-8M | p99 | 0.9747 | 1967 | 2226 | 194 | 1773 |

Continued on next page

**Supplementary Table S11**: Representation-distance-based redundancy reduction summary by percentile. Continued.

| Reduction space | Percentile | Threshold | Retained | Removed | Antioxidant | Non-antioxidant |
|---|---|---|---|---|---|---|
| ESM2-8M | p99.5 | 0.9817 | 2597 | 1596 | 277 | 2320 |
| ESM2-8M | p99.9 | 0.9954 | 3745 | 448 | 584 | 3161 |
| ESMC-300M | p30 | 0.9043 | 34 | 4159 | 4 | 30 |
| ESMC-300M | p40 | 0.9221 | 58 | 4135 | 3 | 55 |
| ESMC-300M | p50 | 0.9346 | 90 | 4103 | 8 | 82 |
| ESMC-300M | p60 | 0.9444 | 124 | 4069 | 12 | 112 |
| ESMC-300M | p70 | 0.9530 | 199 | 3994 | 18 | 181 |
| ESMC-300M | p80 | 0.9612 | 320 | 3873 | 21 | 299 |
| ESMC-300M | p90 | 0.9702 | 588 | 3605 | 38 | 550 |
| ESMC-300M | p95 | 0.9762 | 901 | 3292 | 58 | 843 |
| ESMC-300M | p97 | 0.9796 | 1184 | 3009 | 78 | 1106 |
| ESMC-300M | p98 | 0.9819 | 1456 | 2737 | 91 | 1365 |
| ESMC-300M | p99 | 0.9856 | 2028 | 2165 | 139 | 1889 |
| ESMC-300M | p99.5 | 0.9906 | 2906 | 1287 | 227 | 2679 |
| ESMC-300M | p99.9 | 0.9979 | 3685 | 508 | 514 | 3171 |
| Mistral-Prot | p30 | 0.9976 | 63 | 4130 | 4 | 59 |
| Mistral-Prot | p40 | 0.9979 | 105 | 4088 | 5 | 100 |
| Mistral-Prot | p50 | 0.9982 | 145 | 4048 | 11 | 134 |
| Mistral-Prot | p60 | 0.9984 | 249 | 3944 | 17 | 232 |
| Mistral-Prot | p70 | 0.9986 | 386 | 3807 | 29 | 357 |
| Mistral-Prot | p80 | 0.9988 | 602 | 3591 | 56 | 546 |
| Mistral-Prot | p90 | 0.9991 | 1069 | 3124 | 133 | 936 |
| Mistral-Prot | p95 | 0.9992 | 1549 | 2644 | 242 | 1307 |
| Mistral-Prot | p97 | 0.9993 | 1881 | 2312 | 314 | 1567 |
| Mistral-Prot | p98 | 0.9994 | 2134 | 2059 | 366 | 1768 |
| Mistral-Prot | p99 | 0.9995 | 2518 | 1675 | 445 | 2073 |
| Mistral-Prot | p99.5 | 0.9995 | 2833 | 1360 | 500 | 2333 |
| Mistral-Prot | p99.9 | 0.9996 | 3411 | 782 | 597 | 2814 |
| ProtBERT | p30 | 0.7691 | 24 | 4169 | 3 | 21 |
| ProtBERT | p40 | 0.8088 | 42 | 4151 | 3 | 39 |
| ProtBERT | p50 | 0.8404 | 72 | 4121 | 6 | 66 |
| ProtBERT | p60 | 0.8672 | 107 | 4086 | 12 | 95 |
| ProtBERT | p70 | 0.8916 | 194 | 3999 | 20 | 174 |
| ProtBERT | p80 | 0.9155 | 344 | 3849 | 32 | 312 |
| ProtBERT | p90 | 0.9404 | 709 | 3484 | 72 | 637 |
| ProtBERT | p95 | 0.9561 | 1219 | 2974 | 121 | 1098 |
| ProtBERT | p97 | 0.9642 | 1567 | 2626 | 152 | 1415 |
| ProtBERT | p98 | 0.9692 | 1845 | 2348 | 176 | 1669 |
| ProtBERT | p99 | 0.9760 | 2281 | 1912 | 213 | 2068 |
| ProtBERT | p99.5 | 0.9815 | 2661 | 1532 | 256 | 2405 |
| ProtBERT | p99.9 | 0.9936 | 3537 | 656 | 446 | 3091 |
| ProtT5-XL | p30 | 0.2511 | 16 | 4177 | 2 | 14 |
| ProtT5-XL | p40 | 0.2791 | 22 | 4171 | 2 | 20 |
| ProtT5-XL | p50 | 0.3064 | 30 | 4163 | 2 | 28 |
| ProtT5-XL | p60 | 0.3350 | 37 | 4156 | 3 | 34 |
| ProtT5-XL | p70 | 0.3679 | 65 | 4128 | 5 | 60 |
| ProtT5-XL | p80 | 0.4096 | 111 | 4082 | 7 | 104 |
| ProtT5-XL | p90 | 0.4747 | 224 | 3969 | 17 | 207 |
| ProtT5-XL | p95 | 0.5337 | 469 | 3724 | 23 | 446 |
| ProtT5-XL | p97 | 0.5756 | 685 | 3508 | 37 | 648 |
| ProtT5-XL | p98 | 0.6120 | 914 | 3279 | 45 | 869 |
| ProtT5-XL | p99 | 0.7069 | 1772 | 2421 | 97 | 1675 |
| ProtT5-XL | p99.5 | 0.8888 | 3065 | 1128 | 219 | 2846 |
| ProtT5-XL | p99.9 | 0.9810 | 3678 | 515 | 518 | 3160 |

Continued on next page

**Supplementary Table S11**: Representation-distance-based redundancy reduction summary by percentile. Continued.

| Reduction space | Percentile | Threshold | Retained | Removed | Antioxidant | Non-antioxidant |
|---|---|---|---|---|---|---|
| One-hot | p30–p99.9 | 0.0355–0.2185 | 4193 | 0 | 1010 | 3183 |

#### S4 Splitting strategies and train–test similarity analysis

This section describes how curated and reduced datasets were partitioned for model training and evaluation, and how the resulting train–test separation was assessed. The first subsection summarises the data splitting protocol, including the random, stratified, and distance-aware partitioning strategies used across seeds. The second subsection describes the train–test similarity analysis used to evaluate whether distance-aware splitting produced more stringent evaluation scenarios than conventional random or stratified splits.

##### S4.1 Data splitting protocol

After dataset curation, numerical representation extraction, and redundancy reduction, each dataset version was partitioned into training, validation, and test subsets before model training. Dataset partitioning was performed using `BioSieve`. Three splitting strategies were evaluated: random k-fold splitting, stratified k-fold splitting, and distance-aware k-fold splitting. All splitting procedures were repeated across 30 independent random seeds to quantify the variability associated with data partitioning and to support matched comparisons across experimental configurations.

Each partitioning strategy used five-fold partitioning to generate an 80/20 train–test split, with one fold held out as the test subset. For each split, an additional 10% of the training set was reserved for model selection and validation. This produced approximate final proportions of 72%, 8%, and 20% for training, validation, and test subsets, respectively.

Random k-fold splitting assigned sequences to subsets using shuffled random partitions without explicitly enforcing class proportions. Stratified k-fold splitting used the final binary label to preserve the relative distribution of antioxidant and non-antioxidant examples across train, validation, and test subsets whenever possible. Distance-aware k-fold splitting was used to generate partitions constrained by distances or similarities in a numerical representation space. In this strategy, the representation used to define distances was determined by the active representation-specific configuration. Therefore, distance-aware splits were generated separately for the corresponding representation space rather than using a single fixed representation for all experiments. Ties during distance-aware assignment were shuffled to avoid deterministic ordering effects.

The representation used to construct distance-aware partitions was distinguished from the representation used later as input to the classifier. This distinction allowed the effect of the splitting criterion to be evaluated separately from the effect of the predictive representation. Thus, the same predictive representation could be evaluated under random, stratified, and distance-aware partitions, and distance-aware partitions could be interpreted as representation-space-specific evaluation scenarios.

Sequence identifiers were tracked during partitioning to ensure that no identical sequence appeared in more than one subset within the same split. A split was considered valid only when the required `train.csv`, `val.csv`, and `test.csv` files were generated, no overlap among subsets was detected, at least two classes were present overall, and the generated subsets retained the class diversity required for binary classification and evaluation. Splits were considered invalid when a subset was empty, when train/validation/test overlap was detected, or when the class distribution did not allow model training or performance evaluation. Invalid splits were recorded by the workflow and excluded from downstream model comparison without stopping the complete pipeline.

The main partitioning settings are summarised in Table **S12**.

##### S4.2 Train–test similarity analysis

Train–test similarity was analysed after split generation to quantify how strongly each partitioning strategy separated training and evaluation sequences. This analysis was used as a diagnostic step to compare random, stratified, and distance-aware partitions in terms of the residual similarity between training and test subsets.

**Supplementary Table S12**: Dataset splitting settings used for model training, validation, and testing.

| Setting | Description |
|---|---|
| Partitioning framework | `BioSieve` |
| Splitting strategies | `random_kfold`, `stratified_kfold`, and `distance_aware_kfold` |
| Number of seeds | 30 independent random seeds |
| Number of folds | 5 |
| Initial train/test split | 80% / 20%, derived from five-fold partitioning |
| Validation size | 10% of the training set |
| Approximate final train/validation/test proportions | 72% / 8% / 20% |
| Random split settings | `shuffle = true` |
| Stratified split settings | `shuffle = true`; `dropna = true`; `cast_to_str = false` |
| Distance-aware split settings | `shuffle_ties = true`; distances or similarities defined by the active representation-specific configuration |
| Validation subset | Reserved from the training set and used for model selection and intermediate evaluation |
| Test subset | Held out from model selection and used only for final performance estimation |
| Overlap control | Sequence identifiers were checked to avoid overlap among training, validation, and test subsets |
| Required split files | `train.csv`, `val.csv`, and `test.csv` |
| Invalid split criteria | Splits were excluded if any subset was empty, if overlap was detected, or if the required subsets did not contain sufficient class diversity for binary classification and evaluation |
| Invalid split handling | Invalid splits were recorded without stopping the full workflow |

For each split, the maximum cosine similarity between each test sequence and all sequences in the corresponding training set was computed in the materialised numerical representation space. Feature columns were inferred from the numeric columns shared by the train and test files after excluding identifier, sequence, label, split, seed, and metadata columns. For each test sequence, the highest cosine similarity to any training sequence was retained, and split-level summaries were computed, including the mean and median maximum similarity, the 95th percentile of maximum similarity, the maximum observed similarity, and the proportion of test sequences with maximum similarity greater than or equal to 0.90, 0.95, and 0.99.

The analysis was performed for the random, stratified, and distance-aware splitting strategies across the retained reduction configurations. Distance-based reductions were summarised for the p70, p80, p90, p95, and p99 reduction levels, homology-based reductions for the 0.9, 0.7, 0.5, and 0.3 sequence-identity thresholds, and unreduced datasets as the no-threshold condition. Results were first computed at the fold level and then aggregated by training representation, reduction strategy, reduction level, split representation, and splitting strategy.

Random and stratified splits retained higher train–test similarity than distance-aware splits across all reduction families. This pattern was most pronounced in the unreduced datasets, where the average maximum train–test similarity was 0.81 for both random and stratified splitting, but 0.76 for distance-aware splitting. The proportion of test sequences with maximum similarity greater than or equal to 0.95 also decreased from approximately 25.6% under random or stratified splitting to 12.7% under distance-aware splitting. Similar trends were observed after homology-based and distance-based redundancy reduction, indicating that distance-aware splitting provided an additional separation criterion beyond the reduction procedure itself.

The train–test similarity analysis was interpreted as a validation of the partitioning protocol rather than as an additional dataset construction step. These summaries were used to verify whether distance-aware partitioning produced more stringent evaluation scenarios and to support the interpretation of downstream performance differences across splitting strategies. A summary of the diagnostic analysis is provided in Table **S13**.

#### S5 Machine learning models and hyperparameter grids

This section describes the supervised machine learning models, hyperparameter grids, validation protocol, and performance metrics used to evaluate antioxidant protein prediction across dataset configurations. The modelling stage was designed as a consistent evaluation layer rather than as an optimisation procedure for a single final predictor. Therefore, a predefined set of classical machine learning algorithms and hyperparameter

**Supplementary Table S13**: Train–test similarity diagnostic summary by reduction family and splitting strategy. Values summarise fold-level maximum cosine similarity between each test sequence and its nearest training sequence in the corresponding numerical representation space.

| Reduction family | Split strategy | Mean max. sim. | p95 max. sim. | Test $\geq$ 0.95 | Test $\geq$ 0.99 |
|---|---|---|---|---|---|
| No reduction | Distance-aware | 0.76 | 0.958 | 12.73% | 2.98% |
| No reduction | Random | 0.817 | 0.998 | 25.63% | 12.86% |
| No reduction | Stratified | 0.817 | 0.998 | 25.54% | 12.82% |
| Homology reduction | Distance-aware | 0.737 | 0.911 | 3.23% | 0.16% |
| Homology reduction | Random | 0.771 | 0.948 | 7.88% | 1.00% |
| Homology reduction | Stratified | 0.771 | 0.948 | 7.84% | 1.00% |
| Distance reduction | Distance-aware | 0.652 | 0.806 | 1.17% | 0.11% |
| Distance reduction | Random | 0.676 | 0.842 | 2.88% | 0.47% |
| Distance reduction | Stratified | 0.673 | 0.837 | 2.81% | 0.45% |

configurations was evaluated across the generated combinations of sequence representation, redundancy-control strategy, and splitting strategy.

##### S5.1 Model training and validation protocol

Model training was performed using precomputed external folds generated during the dataset splitting stage. For each experimental configuration, the corresponding `train.csv`, `val.csv`, and `test.csv` files were loaded from the split directory. The training subset was used to fit each model, the validation subset was used for model-selection analyses and intermediate evaluation, and the test subset was kept as an independent held-out set for final performance assessment.

For each algorithm, all predefined hyperparameter configurations were evaluated independently. Each configuration was assigned a configuration index and trained separately on each fold. Predictions were generated for both the validation and test subsets, and the same set of classification metrics was computed for both. The test subset was not used to fit the model or to select hyperparameters. Because the goal of the study was to compare methodological choices across dataset configurations, all evaluated hyperparameter configurations were retained in the output tables rather than using the test set to choose a single best model.

Feature columns were selected from the numerical representation columns available in each split file. The target label was converted to a binary response, with antioxidant proteins encoded as the positive class. Optional feature scaling was supported by the training script; when enabled, the scaler was fitted only on the training subset and then applied to the validation and test subsets through a scikit-learn pipeline. The available scaling options were no scaling, standard scaling, min–max scaling, robust scaling, max-absolute scaling, and $\ell_2$ normalisation. Unless explicitly specified for a given run, models were trained without additional scaling.

Class imbalance was not handled by resampling. Instead, class distribution was controlled at the splitting stage when stratified splitting was used, and model performance was evaluated using metrics that are informative under class imbalance, particularly precision, recall, F1-score, and Matthews correlation coefficient. In the predefined grids, explicit imbalance handling was included only for one XGBoost configuration through `scale_pos_weight = 2`. Model random states were controlled only when specified in the corresponding model configuration. In the predefined grids used here, `random_state = 0` was explicitly set for XGBoost, whereas the experimental seed recorded in the output metadata corresponded to the data partitioning seed.

##### S5.2 Hyperparameter grids

Seven classical machine learning algorithms were evaluated: logistic regression, Gaussian Naive Bayes, k-nearest neighbours, support vector classification, decision tree, random forest, and XGBoost. The predefined hyperparameter configurations used for these algorithms are listed in Table **S14**. These configurations were evaluated as explicit candidate settings rather than as a full Cartesian product of all listed parameter values.

**Supplementary Table S14**: Machine learning algorithms and predefined hyperparameter configurations evaluated in the study. Each row corresponds to one configuration from the hyperparameter grid.

| **Algorithm** | **Cfg.** | **Hyperparameters** |
|---|---|---|
| Logistic Regression | 1 | $C = 0.1$, penalty = `l2`, solver = `lbfgs`, max_iter = 500, n_jobs = 8 |
| Logistic Regression | 2 | $C = 1.0$, penalty = `l2`, solver = `lbfgs`, max_iter = 500, n_jobs = 8 |
| Logistic Regression | 3 | $C = 0.1$, penalty = `l2`, solver = `liblinear`, max_iter = 500, n_jobs = 8 |
| Logistic Regression | 4 | $C = 1.0$, penalty = `l2`, solver = `liblinear`, max_iter = 500, n_jobs = 8 |
| Gaussian Naive Bayes | 1 | var_smoothing = $10^{-9}$ |
| Gaussian Naive Bayes | 2 | var_smoothing = $10^{-8}$ |
| Gaussian Naive Bayes | 3 | var_smoothing = $10^{-7}$ |
| k-nearest neighbours | 1 | n_neighbors = 3, weights = `uniform`, $p = 2$, n_jobs = 8 |
| k-nearest neighbours | 2 | n_neighbors = 5, weights = `uniform`, $p = 2$, n_jobs = 8 |
| k-nearest neighbours | 3 | n_neighbors = 15, weights = `uniform`, $p = 2$, n_jobs = 8 |
| k-nearest neighbours | 4 | n_neighbors = 5, weights = `distance`, $p = 2$, n_jobs = 8 |
| k-nearest neighbours | 5 | n_neighbors = 15, weights = `distance`, $p = 2$, n_jobs = 8 |
| SVC | 1 | $C = 0.1$, kernel = `rbf`, gamma = `scale` |
| SVC | 2 | $C = 1.0$, kernel = `rbf`, gamma = `scale` |
| SVC | 3 | $C = 10$, kernel = `rbf`, gamma = `scale` |
| SVC | 4 | $C = 0.1$, kernel = `linear`, gamma = `scale` |
| SVC | 5 | $C = 1.0$, kernel = `linear`, gamma = `scale` |
| Decision Tree | 1 | max_depth = `None`, min_samples_split = 2, min_samples_leaf = 1, criterion = `gini` |
| Decision Tree | 2 | max_depth = 10, min_samples_split = 2, min_samples_leaf = 2, criterion = `gini` |
| Decision Tree | 3 | max_depth = 30, min_samples_split = 5, min_samples_leaf = 1, criterion = `gini` |
| Decision Tree | 4 | max_depth = 10, min_samples_split = 2, min_samples_leaf = 1, criterion = `entropy` |
| Decision Tree | 5 | max_depth = 30, min_samples_split = 5, min_samples_leaf = 2, criterion = `entropy` |
| Random Forest | 1 | n_estimators = 100, max_depth = `None`, min_samples_split = 2, min_samples_leaf = 1, bootstrap = `true`, n_jobs = 8 |
| Random Forest | 2 | n_estimators = 100, max_depth = 10, min_samples_split = 2, min_samples_leaf = 2, bootstrap = `true`, n_jobs = 8 |
| Random Forest | 3 | n_estimators = 100, max_depth = 30, min_samples_split = 5, min_samples_leaf = 1, bootstrap = `false`, n_jobs = 8 |
| Random Forest | 4 | n_estimators = 300, max_depth = `None`, min_samples_split = 5, min_samples_leaf = 2, bootstrap = `true`, n_jobs = 8 |
| Random Forest | 5 | n_estimators = 300, max_depth = 10, min_samples_split = 2, min_samples_leaf = 1, bootstrap = `false`, n_jobs = 8 |
| Random Forest | 6 | n_estimators = 300, max_depth = 30, min_samples_split = 5, min_samples_leaf = 2, bootstrap = `true`, n_jobs = 8 |
| XGBoost | 1 | n_estimators = 300, max_depth = 4, learning_rate = 0.05, subsample = 0.8, colsample_bytree = 0.8, gamma = 0.0, reg_alpha = 0.0, reg_lambda = 1.0, n_jobs = -1, random_state = 0, eval_metric = `logloss`, use_label_encoder = `false` |
| XGBoost | 2 | n_estimators = 500, max_depth = 6, learning_rate = 0.05, subsample = 0.9, colsample_bytree = 0.9, gamma = 0.0, reg_alpha = 0.0, reg_lambda = 1.0, n_jobs = -1, random_state = 0, eval_metric = `logloss`, use_label_encoder = `false` |
| XGBoost | 3 | n_estimators = 800, max_depth = 8, learning_rate = 0.03, subsample = 0.9, colsample_bytree = 0.9, gamma = 1.0, reg_alpha = 0.1, reg_lambda = 1.0, n_jobs = -1, random_state = 0, eval_metric = `logloss`, use_label_encoder = `false` |
| XGBoost | 4 | n_estimators = 500, max_depth = 6, learning_rate = 0.05, subsample = 0.9, colsample_bytree = 0.9, gamma = 0.0, reg_alpha = 0.0, reg_lambda = 1.0, scale_pos_weight = 2, n_jobs = -1, random_state = 0, eval_metric = `logloss`, use_label_encoder = `false` |

#### S5.3 Performance metrics

Predictive performance was evaluated on both the validation and test subsets using accuracy, precision, recall, F1-score, and Matthews correlation coefficient (MCC). The antioxidant class was treated as the positive class. Let $TP$ denote true positives, $TN$ true negatives, $FP$ false positives, and $FN$ false negatives. The metrics used in this study were defined as follows:

$$\text{Accuracy} = \frac{TP + TN}{TP + TN + FP + FN}. \tag{1}$$

$$\text{Precision} = \frac{TP}{TP + FP}. \tag{2}$$

$$\text{Recall} = \frac{TP}{TP + FN}. \tag{3}$$

$$\text{F1} = \frac{2TP}{2TP + FP + FN}. \tag{4}$$

$$\text{MCC} = \frac{(TP \times TN) - (FP \times FN)}{\sqrt{(TP + FP)(TP + FN)(TN + FP)(TN + FN)}}. \tag{5}$$

Precision, recall, and F1-score were computed with undefined divisions set to zero. This avoids undefined metric values in cases where a model predicts no positive examples or when a class is absent from the prediction output. MCC was used as the primary performance metric because it incorporates all four entries of the confusion matrix and is informative under class imbalance. F1-score was used as a complementary metric, whereas accuracy, precision, and recall were treated as secondary metrics.

##### S5.4 Aggregation of model performance

For each model, hyperparameter configuration, seed, partitioning strategy, representation, redundancy strategy, and fold, performance metrics were recorded separately for the validation and test subsets. Fold-level outputs included the number of training, validation, and test examples, the number of positive and negative examples in each subset, the number of numerical features, the scaler used, the hyperparameter configuration, and the validation and test performance metrics.

Fold-level results were subsequently aggregated for each experimental configuration. Aggregation was performed by grouping results according to the model, hyperparameter configuration, seed, representation, redundancy-control strategy, splitting strategy, reduction level, and split space when applicable. This structure preserved the distinction between the representation used for model training, the representation or criterion used for redundancy reduction, and the representation space used for distance-aware splitting.

For each metric, the mean, standard deviation, and number of contributing folds were computed separately for validation and test performance. These summaries were used to compare model behaviour across sequence representations, redundancy-control strategies, and splitting scenarios. Invalid or failed model fits were recorded during training and excluded from aggregate summaries.

For the main aggregation tables, model performance was reported as mean and standard deviation across valid folds and seeds. Confidence intervals were not used as the default summary statistic for the complete set of model outputs.

For selected downstream summaries of representation-role sensitivity configurations reported under distance-aware partitioning, 95% confidence intervals were computed explicitly from the aggregated test-set MCC values. For a given configuration, the 95% confidence interval was calculated as

$$\bar{x} \pm 1.96 \times \frac{s}{\sqrt{n}},$$

where $\bar{x}$ is the mean test MCC, $s$ is the standard deviation of the contributing seed-level aggregated MCC values, and $n$ is the number of valid seed-level aggregated observations used for that configuration. In these summaries, configurations were retained only when at least 30 unique matched seeds were available. The number of contributing rows and the number of unique seeds were recorded to support reproducibility of the reported intervals.

#### S6 Paired methodological delta analyses

This section describes the operational procedure used to compute the paired methodological delta analyses reported in the main text. Metric definitions are provided in Equations 1–5; therefore, this section focuses on how candidate and reference configurations were paired, which baseline was used for each methodological comparison, and how delta values were summarised.

The purpose of the delta analyses was to quantify changes in predictive performance associated with specific methodological decisions, rather than to compare configurations only in terms of absolute performance. For this reason, each candidate configuration was compared against a matched reference configuration. Whenever possible, candidate and reference configurations were matched so that they differed in the methodological factor under analysis while retaining the same representation, model, hyperparameter configuration, seed, reduction level, and splitting context.

##### S6.1 Operational definition of paired deltas

Methodological deltas were computed as candidate performance minus matched reference performance. This convention was applied to each aggregated test-set metric. Because Matthews correlation coefficient (MCC) was the main metric used to visualise and interpret the paired delta analyses in the main text, its operational definition is shown explicitly here:

$$\Delta\text{MCC}(c, r) = \text{MCC}(c) - \text{MCC}(r), \tag{6}$$

where $c$ denotes the candidate configuration and $r$ denotes the matched reference configuration. Positive $\Delta$MCC values indicate higher MCC under the candidate condition, values close to zero indicate that MCC was approximately retained, and negative values indicate lower MCC relative to the matched reference. The same candidate-minus-reference convention was applied to the remaining test-set metrics.

For complementary ranking and interpretability, metric loss was defined as the inverse difference between the matched reference and candidate configurations. For a generic metric $M$, this loss was calculated as

$$\text{Loss}_M(c, r) = M(r) - M(c). \tag{7}$$

Thus, lower loss values indicate better preservation of the reference performance, whereas higher loss values indicate larger performance decreases relative to the matched reference. Because loss is the inverse of the candidate-minus-reference delta, positive loss values correspond to negative deltas and therefore indicate candidate underperformance relative to the reference.

##### S6.2 Reference configurations used for paired comparisons

Each methodological comparison used a reference configuration selected to isolate the effect of the factor under analysis. The reference configuration was not intended to represent the best-performing model; instead, it served as the matched baseline against which a specific methodological change was evaluated. Candidate and reference configurations were paired by matching the remaining experimental factors, including model, hyperparameter configuration, seed, representation, reduction level, and splitting context. The main reference configurations are summarised in Table **S15**.

For partitioning analyses, random k-fold splitting was used as the reference because it represents the conventional baseline partitioning strategy. Stratified and distance-aware partitions were therefore interpreted relative to matched random partitions. For redundancy-control analyses, the unreduced dataset was used as the reference to quantify the performance change associated with applying homology-based or representation-distance-based reduction.

For reduction-space analyses, candidate and reference configurations were matched at the same reduction level whenever possible, but differed in the representation space used to define redundancy. This allowed the effect of the reduction space to be evaluated separately from the effect of reduction intensity. For representation-role analyses, the reference depended on the specific matched comparison, while preserving the distinction between the representation used for model training, the representation used to define redundancy, and the representation space used for distance-aware partitioning.

**Supplementary Table S15**: Reference configurations used for paired methodological delta analyses. Candidate configurations were compared against the corresponding reference while keeping the remaining experimental factors matched whenever possible.

| Analysis | Reference configuration | Candidate configuration | Delta interpretation |
|---|---|---|---|
| Partitioning strategy | Random k-fold split | Stratified k-fold or distance-aware k-fold split | Effect of changing the train–test partitioning rule |
| Redundancy control | No-reduction dataset | Homology-reduced or representation-distance-reduced dataset | Effect of applying redundancy reduction |
| Reduction space | Reduction defined in the matched reference space | Reduction defined at the same reduction level in an alternative representation space | Effect of the representation space used to define redundancy |
| Representation role | Matched reference representation under the same workflow condition | Alternative predictive representation or alternative representation-space condition | Effect of representation choice while preserving the surrounding workflow context |

Pairs were included only when both the candidate and the matched reference configuration were available and valid. Candidate configurations without a valid matched reference were excluded from the corresponding delta analysis.

##### S6.3 Paired matching rules

Candidate–reference pairs were constructed only between configurations that were comparable for the methodological question being evaluated. Matching was performed using the experimental identifiers relevant to each analysis, so that the candidate and reference differed primarily in the methodological factor of interest. The main matching rules are summarised in Table **S16**.

**Supplementary Table S16**: Matching rules used to define valid candidate–reference pairs in the methodological delta analyses. Variables listed as matched were held constant whenever possible so that deltas primarily reflected the factor under analysis.

| Analysis | Variables matched between candidate and reference configurations |
|---|---|
| Partitioning strategy | Training representation, classifier, scaler, hyperparameter configuration, random seed, redundancy-control strategy, reduction label, reduction space, reduction level, reduction percentile, and homology threshold |
| Redundancy control | Training representation, classifier, partitioning strategy, scaler, hyperparameter configuration, and random seed |
| Reduction space | Training representation, classifier, partitioning strategy, redundancy-control family, reduction level or threshold, scaler, hyperparameter configuration, and random seed, whenever applicable |
| Representation role | Classifier, partitioning strategy, redundancy-control condition, scaler, hyperparameter configuration, random seed, and the corresponding matched dataset condition defined for the comparison |

This matching scheme was designed to avoid comparing configurations that differed simultaneously in multiple uncontrolled factors. For example, when evaluating the effect of a partitioning strategy, the classifier, training representation, seed, scaler, hyperparameter configuration, and redundancy-control condition were kept fixed whenever the corresponding matched reference was available. Similarly, when evaluating redundancy control, the partitioning strategy and modelling configuration were held constant, and the reduced condition was compared against the corresponding unreduced reference.

##### S6.4 Validity criteria for paired comparisons

Paired delta values were calculated only for candidate–reference pairs that were available after dataset generation, split validation, model training, and metric aggregation. A candidate–reference pair was considered valid when all of the following criteria were met:

- Both the candidate and reference configurations completed model fitting successfully.
- Both configurations had the required aggregated test-set metric available.

- Candidate and reference configurations could be matched according to the comparison-specific rules described above.
- The matched pair differed in the methodological factor under analysis while preserving the remaining experimental factors whenever possible.
- The paired comparison corresponded to the same metric, model, hyperparameter configuration, seed, and relevant workflow context.

Configurations were excluded from a paired delta analysis when no corresponding matched reference was available. As a result, the number of valid pairs was allowed to differ across methodological comparisons, metrics, representations, reduction strategies, reduction levels, classifiers, and partitioning strategies. This avoided estimating deltas from non-equivalent configurations and ensured that each delta represented a within-setting change relative to a comparable baseline.

Invalid splits or failed model fits were handled upstream during model training and aggregation. The paired delta analysis operated on the aggregated performance table and therefore used only completed candidate and reference results available for valid paired comparison.

##### S6.5 Aggregation of delta values

For each methodological comparison, deltas were summarised across random seeds and other valid matched settings. The average delta for a metric $M$ was calculated as

$$\overline{\Delta M} = \frac{1}{N}\sum_{i=1}^{N}\left[M(c_i) - M(r_i)\right], \tag{8}$$

where $N$ is the number of valid matched candidate–reference pairs. In addition to the mean delta, summaries included the median delta, the standard deviation of delta values, the number of valid pairs, and the number of random seeds represented in each grouped comparison.

Performance retention was also computed as

$$\text{Retention}_M(c, r) = \frac{M(c)}{M(r)}, \tag{9}$$

when the reference metric value was non-zero. Retention values close to one indicate that the candidate configuration preserved the reference performance, whereas lower values indicate stronger performance loss.

For ranking summaries, configurations were ordered primarily by MCC loss relative to the matched reference, with lower MCC loss indicating better performance preservation. F1-score was used as a complementary ranking metric, whereas accuracy, precision, and recall were treated as secondary metrics. Configurations with insufficient paired evidence were retained in the output summaries but were not interpreted as recommended configurations.

#### S7 Sensitivity analyses

This section summarises the sensitivity analyses performed to assess whether the main conclusions were robust to selected changes in dataset-support criteria, similarity-control strength, and representation roles within the workflow. These analyses were not intended to exhaustively evaluate all possible combinations of methodological choices. Instead, they provided targeted checks of factors that could influence dataset composition, evaluation difficulty, and apparent predictive performance.

##### S7.1 Sensitivity analysis design

Four sensitivity analyses were organised into three methodological groups. First, source-support analyses evaluated whether different levels of concordant source-entry support altered dataset composition and downstream model performance. Second, redundancy and similarity degradation analyses evaluated how predictive performance changed as sequence-identity or representation-distance constraints became more restrictive. Third, representation-role analyses evaluated whether the representation used to define redundancy control or distance-aware partitioning influenced performance beyond its role as model input.

The sensitivity analyses are summarised in Table **S17**. Model evaluation followed the same splitting, training, metric-calculation, and aggregation procedures described in Sections S4 and S5. Performance changes were interpreted using the paired delta framework described in Section S6.

**Supplementary Table S17**: Overview of sensitivity analyses performed to evaluate the robustness of the main methodological conclusions.

| Analysis | Variable tested | Purpose |
|---|---|---|
| Source-entry support | Number of source-level dataset entries providing concordant labels | Assess how different levels of documentary concordance are associated with dataset composition and model performance |
| Homology degradation | Sequence identity threshold | Assess the effect of increasingly stringent homology-based redundancy control |
| Representation-distance degradation | Representation-specific similarity or distance threshold | Assess the effect of increasingly stringent representation-based similarity control |
| Representation-role analysis | Predictive, reduction, and split representation spaces | Assess whether the representation used to structure the dataset affects performance beyond its role as model input |

##### S7.2 Source-support sensitivity analysis

The source-support analysis evaluated whether the amount of concordant documentary reporting associated with each retained sequence influenced dataset composition and downstream predictive performance. Sequence-level subsets were constructed from the final harmonised dataset using the number of source-level dataset entries providing concordant labels for each sequence. The same supervised-learning workflow used for the main analyses was then applied to each source-support condition, so that the modified factor was the input dataset subset.

The source-support conditions should be interpreted according to their operational definitions rather than as four mutually exclusive groups or as levels of experimental validation. The single-source and multi-source conditions separate sequences reported by exactly one source-level dataset entry from sequences reported concordantly by at least two entries. The high-support condition is a stricter subset of the multi-source condition, whereas the complete-consensus condition corresponds to the full curated dataset used in the main analyses after harmonisation, conflict resolution, and filtering. The terms single-source, multi-source, and high-support are retained as operational subset names and do not imply that the contributing entries represent independent experiments.

Definitions and dataset composition values for the source-support sensitivity analysis are summarised in Table **S18**. These values complement the visual summary presented in the main text by reporting the sequence counts and class proportions used in the analysis.

**Supplementary Table S18**: Source-support conditions and dataset composition used in the source-support sensitivity analysis. Positive sequences correspond to antioxidant proteins and negative sequences correspond to non-antioxidant proteins.

| Condition | Definition | Total | Positive | Negative | Positive (%) |
|---|---|---|---|---|---|
| Single-source | $n_{\text{valid source entries}} = 1$ | 1895 | 573 | 1322 | 30.2 |
| Multi-source | $n_{\text{valid source entries}} \geq 2$ | 2298 | 437 | 1861 | 19.0 |
| High-support | $n_{\text{valid source entries}} \geq 3$ | 2207 | 347 | 1860 | 15.7 |
| Complete consensus | All retained sequences | 4193 | 1010 | 3183 | 24.1 |

To support the source-support performance trends shown in main-text Figure 2D, Table **S19** reports the MCC values used for the plotted source-support subsets. These values were computed directly from the source-support training results. For each source-support condition and partitioning strategy, test-set MCC values were aggregated across valid representation–algorithm–seed evaluations. The resulting summaries report the mean MCC, the corresponding 95% confidence interval, the number of evaluated entries, and the number of unique seeds.

**Supplementary Table S19**: Source-support sensitivity performance values used in main-text Figure 2D. Values correspond to mean test-set MCC values computed directly from valid representation–algorithm–seed evaluations.

| Condition | Split strategy | Mean MCC | 95% CI | Evaluated entries | Unique seeds |
|---|---|---|---|---|---|
| Single-source | Random | 0.941 | 0.0042 | 270 | 30 |
| Multi-source | Random | 0.777 | 0.0060 | 270 | 30 |
| High-support | Random | 0.764 | 0.0052 | 270 | 30 |
| Complete consensus | Random | 0.882 | 0.0034 | 540 | 30 |
| Single-source | Stratified | 0.940 | 0.0042 | 270 | 30 |
| Multi-source | Stratified | 0.778 | 0.0059 | 270 | 30 |
| High-support | Stratified | 0.762 | 0.0051 | 270 | 30 |
| Complete consensus | Stratified | 0.882 | 0.0035 | 540 | 30 |
| Single-source | Distance-aware | 0.821 | 0.0052 | 270 | 30 |
| Multi-source | Distance-aware | 0.668 | 0.0083 | 270 | 30 |
| High-support | Distance-aware | 0.644 | 0.0101 | 270 | 30 |
| Complete consensus | Distance-aware | 0.724 | 0.0046 | 540 | 30 |

*Note.* Values correspond to the MCC points displayed in main-text Figure 2D. Mean MCC values were computed directly from the source-support training results rather than reconstructed from the training-trend ranking summary. Error estimates correspond to 95% confidence intervals computed across valid representation–algorithm–seed evaluations. Evaluated entries correspond to the valid rows available after source-support training and filtering for each condition and partitioning strategy. Random and stratified partitioning produced nearly identical trends, but they are reported separately because they were evaluated as distinct partitioning strategies.

##### S7.3 Similarity-control sensitivity analysis

The similarity-control sensitivity analysis evaluated whether progressively stricter redundancy or similarity constraints altered downstream model performance. Two complementary forms of similarity control were considered: homology-based degradation and representation-distance-based degradation. Both analyses reused the datasets generated during the redundancy-reduction workflow described in Section S3.

The sensitivity design for similarity-control analyses is summarised in Table **S20**. In both cases, performance changes were summarised as paired deltas relative to the corresponding no-reduction reference condition. This design allowed the effect of increasing similarity control to be evaluated while minimising comparisons between non-equivalent experimental settings.

**Supplementary Table S20**: Design of the homology-based and representation-distance-based similarity-control sensitivity analyses.

| Analysis | Reference condition | Candidate conditions | Interpretation |
|---|---|---|---|
| Homology degradation | Unreduced dataset | 0.9, 0.7, 0.5, and 0.3 minimum sequence identity | Effect of increasingly stringent sequence-identity-based redundancy control |
| Representation-distance degradation | Unreduced dataset | Representation-specific percentile thresholds from p99.9 to p40, depending on split validity and available matched comparisons | Effect of increasingly stringent representation-space similarity control |

For homology-based degradation, the unreduced dataset was compared with datasets reduced at 0.9, 0.7, 0.5, and 0.3 minimum sequence identity. Lower identity thresholds correspond to stricter homology control because they collapse sequences at lower levels of sequence similarity. To support the homology-control component of the similarity-control analysis, Table **S21** reports test-set MCC values after homology-based redundancy reduction. For each identity threshold and partitioning strategy, performance was compared against the matched no-reduction reference. Pairing was performed by matching the model-input representation, split space, partitioning strategy, classifier, scaler, hyperparameter configuration, and random seed. Therefore, the reported ΔMCC values quantify the change in performance associated with homology-based redundancy reduction under otherwise matched evaluation settings.

**Supplementary Table S21**: Homology-control degradation performance summary. Values report test-set MCC after homology-based redundancy reduction and paired changes relative to the matched no-reduction reference condition.

| Identity threshold | Retained sequences | Split strategy | No-reduction MCC | Homology-reduced MCC | Mean ΔMCC | Valid pairs |
|---|---|---|---|---|---|---|
| No reduction | 4193 | Random | 0.702 | 0.702 | 0.000 | 12480 |
| No reduction | 4193 | Stratified | 0.702 | 0.702 | 0.000 | 12480 |
| No reduction | 4193 | Distance-aware | 0.562 | 0.562 | 0.000 | 11520 |
| 0.9 | 3801 | Random | 0.702 | 0.597 | -0.106 | 12480 |
| 0.9 | 3801 | Stratified | 0.702 | 0.597 | -0.106 | 12480 |
| 0.9 | 3801 | Distance-aware | 0.562 | 0.493 | -0.069 | 11520 |
| 0.7 | 3510 | Random | 0.702 | 0.434 | -0.268 | 12480 |
| 0.7 | 3510 | Stratified | 0.702 | 0.433 | -0.269 | 12480 |
| 0.7 | 3510 | Distance-aware | 0.562 | 0.351 | -0.210 | 11520 |
| 0.5 | 3252 | Random | 0.702 | 0.271 | -0.431 | 12480 |
| 0.5 | 3252 | Stratified | 0.702 | 0.271 | -0.431 | 12480 |
| 0.5 | 3252 | Distance-aware | 0.562 | 0.222 | -0.340 | 11520 |
| 0.3 | 2948 | Random | 0.702 | 0.143 | -0.559 | 12480 |
| 0.3 | 2948 | Stratified | 0.702 | 0.144 | -0.559 | 12480 |
| 0.3 | 2948 | Distance-aware | 0.562 | 0.121 | -0.441 | 11520 |

*Note.* ΔMCC was computed as homology-reduced MCC minus the matched no-reduction MCC. The no-reduction MCC column corresponds to the matched reference used for each paired comparison, not to an independent global average. Values were summarised across valid matched representation–split-space–classifier–scaler–configuration–seed combinations. Distance-aware rows include the standard distance-aware partitioning workflow; one-hot-specific normalised and non-normalised distance-aware variants were not pooled into this summary.

For representation-distance-based degradation, the unreduced dataset was compared with representation-specific percentile-based reductions. Because these percentiles are computed within each representation-specific similarity or distance distribution, they were interpreted within each reduction space rather than as universal similarity thresholds shared across embeddings. Dataset sizes, retained sequence counts, and class distributions for the corresponding representation-distance-based reductions are reported in Section S3.

To support the representation-distance component of the similarity-control analysis, Table **S22** reports paired changes in test-set MCC and F1-score after representation-distance-based redundancy reduction. Each reduced configuration was compared against the matched no-reduction reference by matching the model-input representation, partitioning strategy, classifier, scaler, hyperparameter configuration, and random seed. Therefore, the reported ΔMCC and ΔF1 values quantify the performance change associated with representation-distance-based reduction under otherwise matched evaluation settings.

**Supplementary Table S22**: Representation-distance degradation performance summary. Values report paired changes in test-set MCC and F1-score relative to the matched no-reduction reference condition.

| Reduction space | Threshold | Split strategy | No-reduction MCC | Reduced MCC | ΔMCC | ΔF1 | Valid seeds |
|---|---|---|---|---|---|---|---|
| Ankh2-ext1 | p99.9 | Distance-aware | 0.575 | 0.601 | 0.026 | 0.013 | 30 |
| Ankh2-ext1 | p99.5 | Distance-aware | 0.575 | 0.317 | -0.258 | -0.296 | 30 |
| Ankh2-ext1 | p99 | Distance-aware | 0.575 | 0.124 | -0.451 | -0.480 | 30 |
| Ankh2-ext1 | p98 | Distance-aware | 0.576 | 0.066 | -0.510 | -0.524 | 27 |
| Ankh2-ext1 | p97 | Distance-aware | 0.575 | 0.048 | -0.527 | -0.542 | 27 |
| Ankh2-ext1 | p95 | Distance-aware | 0.576 | 0.031 | -0.546 | -0.555 | 18 |
| Ankh2-ext1 | p90 | Distance-aware | 0.576 | -0.003 | -0.579 | -0.586 | 11 |
| Ankh2-ext1 | p80 | Distance-aware | 0.571 | 0.002 | -0.569 | -0.573 | 6 |
| ESM2-8M | p99.9 | Distance-aware | 0.630 | 0.550 | -0.080 | -0.094 | 30 |
| ESM2-8M | p99.5 | Distance-aware | 0.630 | 0.478 | -0.152 | -0.177 | 30 |
| ESM2-8M | p99 | Distance-aware | 0.630 | 0.448 | -0.182 | -0.202 | 30 |
| ESM2-8M | p98 | Distance-aware | 0.630 | 0.435 | -0.196 | -0.218 | 30 |
| ESM2-8M | p97 | Distance-aware | 0.630 | 0.422 | -0.208 | -0.231 | 30 |
| ESM2-8M | p95 | Distance-aware | 0.630 | 0.317 | -0.313 | -0.335 | 30 |
| ESM2-8M | p90 | Distance-aware | 0.630 | 0.220 | -0.410 | -0.431 | 30 |
| ESM2-8M | p80 | Distance-aware | 0.631 | 0.147 | -0.483 | -0.497 | 20 |
| ESM2-8M | p70 | Distance-aware | 0.631 | 0.014 | -0.617 | -0.616 | 14 |

*Continued on the next page*

| Reduction space | Threshold | Split strategy | No-reduction MCC | Reduced MCC | ΔMCC | ΔF1 | Valid seeds |
|---|---|---|---|---|---|---|---|
| ESM2-8M | p60 | Distance-aware | 0.631 | 0.022 | -0.609 | -0.611 | 14 |
| ESM2-8M | p50 | Distance-aware | 0.631 | 0.011 | -0.619 | -0.625 | 4 |
| ESM2-8M | p40 | Distance-aware | 0.630 | 0.010 | -0.620 | -0.627 | 2 |
| ESMC-300M | p99.9 | Distance-aware | 0.553 | 0.438 | -0.115 | -0.136 | 30 |
| ESMC-300M | p99.5 | Distance-aware | 0.553 | 0.191 | -0.362 | -0.422 | 30 |
| ESMC-300M | p99 | Distance-aware | 0.553 | 0.105 | -0.447 | -0.490 | 30 |
| ESMC-300M | p98 | Distance-aware | 0.553 | 0.053 | -0.499 | -0.528 | 30 |
| ESMC-300M | p97 | Distance-aware | 0.552 | 0.061 | -0.491 | -0.522 | 29 |
| ESMC-300M | p95 | Distance-aware | 0.553 | 0.037 | -0.516 | -0.541 | 29 |
| ESMC-300M | p90 | Distance-aware | 0.552 | 0.045 | -0.507 | -0.533 | 26 |
| ESMC-300M | p80 | Distance-aware | 0.554 | 0.010 | -0.544 | -0.567 | 11 |
| ESMC-300M | p70 | Distance-aware | 0.552 | 0.016 | -0.536 | -0.550 | 8 |
| ESMC-300M | p60 | Distance-aware | 0.553 | 0.002 | -0.550 | -0.571 | 3 |
| Mistral-Prot | p99.9 | Distance-aware | 0.343 | 0.236 | -0.107 | -0.142 | 30 |
| Mistral-Prot | p99.5 | Distance-aware | 0.343 | 0.238 | -0.105 | -0.145 | 30 |
| Mistral-Prot | p99 | Distance-aware | 0.343 | 0.225 | -0.118 | -0.155 | 30 |
| Mistral-Prot | p98 | Distance-aware | 0.343 | 0.212 | -0.131 | -0.170 | 30 |
| Mistral-Prot | p97 | Distance-aware | 0.343 | 0.203 | -0.141 | -0.184 | 30 |
| Mistral-Prot | p95 | Distance-aware | 0.343 | 0.195 | -0.148 | -0.190 | 30 |
| Mistral-Prot | p90 | Distance-aware | 0.343 | 0.129 | -0.214 | -0.263 | 30 |
| Mistral-Prot | p80 | Distance-aware | 0.343 | 0.059 | -0.285 | -0.329 | 29 |
| Mistral-Prot | p70 | Distance-aware | 0.343 | 0.044 | -0.299 | -0.348 | 22 |
| ProtBERT | p99.9 | Distance-aware | 0.696 | 0.490 | -0.205 | -0.243 | 30 |
| ProtBERT | p99.5 | Distance-aware | 0.696 | 0.320 | -0.376 | -0.442 | 30 |
| ProtBERT | p99 | Distance-aware | 0.696 | 0.273 | -0.423 | -0.490 | 30 |
| ProtBERT | p98 | Distance-aware | 0.696 | 0.246 | -0.449 | -0.514 | 30 |
| ProtBERT | p97 | Distance-aware | 0.696 | 0.241 | -0.455 | -0.516 | 30 |
| ProtBERT | p95 | Distance-aware | 0.696 | 0.238 | -0.458 | -0.517 | 30 |
| ProtBERT | p90 | Distance-aware | 0.696 | 0.171 | -0.525 | -0.571 | 30 |
| ProtBERT | p80 | Distance-aware | 0.696 | 0.091 | -0.605 | -0.642 | 23 |
| ProtBERT | p70 | Distance-aware | 0.696 | 0.071 | -0.625 | -0.658 | 15 |
| ProtBERT | p60 | Distance-aware | 0.696 | 0.033 | -0.663 | -0.690 | 4 |
| ProtT5-XL | p99.9 | Distance-aware | 0.582 | 0.530 | -0.052 | -0.077 | 30 |
| ProtT5-XL | p99.5 | Distance-aware | 0.582 | 0.297 | -0.285 | -0.325 | 30 |
| ProtT5-XL | p99 | Distance-aware | 0.582 | 0.115 | -0.467 | -0.496 | 30 |
| ProtT5-XL | p98 | Distance-aware | 0.582 | 0.086 | -0.496 | -0.518 | 24 |
| ProtT5-XL | p97 | Distance-aware | 0.582 | 0.039 | -0.543 | -0.554 | 26 |
| ProtT5-XL | p95 | Distance-aware | 0.581 | 0.006 | -0.575 | -0.585 | 15 |
| ProtT5-XL | p90 | Distance-aware | 0.581 | 0.020 | -0.561 | -0.570 | 6 |

*Note.* ΔMCC and ΔF1 were computed as the reduced-condition performance minus the matched no-reduction reference. Pairing was performed by matching model-input representation, partitioning strategy, classifier, scaler, hyperparameter configuration, and random seed. Valid seeds indicate the number of unique seeds contributing to each matched summary after filtering.

Overall, the representation-distance degradation analysis showed that increasingly restrictive percentile thresholds generally reduced predictive performance relative to the matched no-reduction reference. The magnitude of the decrease differed across representation spaces, supporting the interpretation that representation-aware redundancy control changes not only the amount of retained data, but also downstream evaluation difficulty and the number of valid matched comparisons available for performance estimation.

Because increasingly restrictive thresholds can reduce dataset size, class diversity, and the number of valid train–validation–test partitions, Table **S23** summarises representative cases in which fewer than 30 seeds remained available after filtering. This table is intended to document the feasibility effect of strict similarity control rather than to provide an exhaustive list of all invalid configurations.

##### S7.4 Representation-role sensitivity analysis

The representation-role sensitivity analysis evaluated whether the representation used to define dataset structure influenced model performance beyond its use as input features for the classifier. In the main representation-specific workflow, representation roles were matched whenever possible. Configurations in which the training representation, reduction space, or split space differed were analysed separately as representation-role sensitivity analyses.

**Supplementary Table S23**: Validity summary for selected strict representation-distance configurations.

| Reduction space | Threshold | Attempted seeds | Valid seeds | Filtered seeds | Interpretation |
|---|---|---|---|---|---|
| Ankh2-ext1 | p90 | 30 | 11 | 19 | Reduced split validity under stringent filtering |
| Ankh2-ext1 | p80 | 30 | 6 | 24 | Reduced split validity under stringent filtering |
| ESM2-8M | p50 | 30 | 4 | 26 | Reduced split validity under stringent filtering |
| ESM2-8M | p40 | 30 | 2 | 28 | Reduced split validity under stringent filtering |
| ESMC-300M | p70 | 30 | 8 | 22 | Reduced split validity under stringent filtering |
| ESMC-300M | p60 | 30 | 3 | 27 | Reduced split validity under stringent filtering |
| ProtBERT | p70 | 30 | 15 | 15 | Reduced split validity under stringent filtering |
| ProtBERT | p60 | 30 | 4 | 26 | Reduced split validity under stringent filtering |
| ProtT5-XL | p95 | 30 | 15 | 15 | Reduced split validity under stringent filtering |
| ProtT5-XL | p90 | 30 | 6 | 24 | Reduced split validity under stringent filtering |

*Note.* Attempted seeds correspond to the planned number of random seeds for each configuration. Valid seeds correspond to the number of seeds retained in the matched summary after filtering for valid splits, complete model outputs, and available candidate–reference pairs. Filtered seeds were not necessarily failed model runs; they include seeds excluded because the corresponding reduced dataset, split, or matched comparison was not valid for paired performance analysis.

Two targeted designs were evaluated. In Design 1, the training representation was fixed and the representation-distance reduction space was varied. In this workflow, distance-aware splitting was computed using the training representation, so the split space remained matched to the training representation. In Design 2, the reduction and split spaces were fixed to a common representation space while the training representation was varied.

The representation-role configurations are summarised in Table **S24**. These analyses were based on targeted configurations rather than on a full factorial crossing of all possible training, reduction, and split representation spaces.

**Supplementary Table S24**: Representation-role configurations evaluated in the sensitivity analysis. Design 1 fixed the training and split spaces while varying the reduction space. Design 2 fixed the reduction and split spaces while varying the training representation.

| Design | Condition | Reduction space | Split space | Training representation |
|---|---|---|---|---|
| Design 1 | Ankh2-ext1 fixed | Ankh2-ext1, ESM2-8M, ESMC-300M, Mistral-Prot, ProtBERT, ProtT5-XL | Ankh2-ext1 | Ankh2-ext1 |
| Design 1 | ESM2-8M fixed | ESM2-8M, Ankh2-ext1, ESMC-300M, Mistral-Prot, ProtBERT, ProtT5-XL | ESM2-8M | ESM2-8M |
| Design 1 | ESMC-300M fixed | ESMC-300M, Ankh2-ext1, ESM2-8M, Mistral-Prot, ProtBERT, ProtT5-XL | ESMC-300M | ESMC-300M |
| Design 1 | Mistral-Prot fixed | Mistral-Prot, Ankh2-ext1, ESM2-8M, ESMC-300M, ProtBERT, ProtT5-XL | Mistral-Prot | Mistral-Prot |
| Design 1 | ProtBERT fixed | ProtBERT, Ankh2-ext1, ESM2-8M, ESMC-300M, Mistral-Prot, ProtT5-XL | ProtBERT | ProtBERT |
| Design 1 | ProtT5-XL fixed | ProtT5-XL, Ankh2-ext1, ESM2-8M, ESMC-300M, Mistral-Prot, ProtBERT | ProtT5-XL | ProtT5-XL |
| Design 1 | One-hot fixed | Ankh2-ext1, ESM2-8M, ESMC-300M, Mistral-Prot, ProtBERT, ProtT5-XL | One-hot | One-hot |
| Design 2 | Ankh2-ext1 train, Mistral-Prot-structured | Mistral-Prot | Mistral-Prot | Ankh2-ext1 |
| Design 2 | ProtT5-XL train, Mistral-Prot-structured | Mistral-Prot | Mistral-Prot | ProtT5-XL |
| Design 2 | Mistral-Prot train, Mistral-Prot-structured | Mistral-Prot | Mistral-Prot | Mistral-Prot |

No-reduction and homology-reduction configurations were retained as reference or control conditions, but were not interpreted as off-diagonal representation-distance configurations because redundancy was not defined through an alternative embedding space. When paired or delta-based comparisons were computed, only valid matched configurations were used, as described in Section S6.

To support the role-decoupled configurations reported in the main text, Table **??** summarises selected representation-role configurations observed in the targeted representation-role sensitivity analysis under distance-aware partitioning. The configurations are ordered descriptively by mean test-set MCC and correspond to cases in which the model-input representation differed from the representation space used for redundancy reduction and distance-aware partitioning. They are reported to illustrate the outcomes of the targeted sensitivity analysis and do not represent the selection of a final antioxidant protein predictor.

| Rank | Input rep. | Reduction setting | Split space | Algorithm | MCC mean | 95% CI | ΔMCC | Valid seeds |
|---|---|---|---|---|---|---|---|---|
| 1 | ProtT5-XL | Mistral-Prot p90 | Mistral-Prot | SVC cfg. 2 | 0.84 | 0.003 | -0.005 | 30 |
| 2 | Ankh2-ext1 | Mistral-Prot p90 | Mistral-Prot | SVC cfg. 2 | 0.82 | 0.004 | -0.02 | 30 |
| 3 | Ankh2-ext1 | Mistral-Prot p90 | Mistral-Prot | XGBoost cfg. 3 | 0.79 | 0.004 | -0.008 | 30 |
| 4 | Ankh2-ext1 | Mistral-Prot p90 | Mistral-Prot | k-NN cfg. 4 | 0.75 | 0.007 | -0.02 | 30 |
| 5 | ProtT5-XL | Mistral-Prot p90 | Mistral-Prot | XGBoost cfg. 3 | 0.74 | 0.006 | -0.03 | 30 |

*Note.* MCC values are reported as mean and 95% confidence interval computed from valid seed-level aggregated test-set MCC values, as described in Section S5.4. Only configurations with at least 30 unique matched seeds were retained. ΔMCC corresponds to the mean difference relative to the matched random-split reference. The input representation was used as the supervised-learning feature space. The reduction setting reports the representation space and percentile threshold used for representation-distance-based filtering. The split space reports the representation used to compute train–test proximity during distance-aware partitioning. These configurations are reported as representation-role sensitivity outcomes and should not be interpreted as part of the matched representation-specific workflow. They should not be interpreted as a final model selection or as recommended predictor configurations.

#### S8 Implementation details, software versions, and reproducibility

This section summarises the computational environment and reproducibility records used to generate the curated datasets, numerical representations, reduced datasets, data splits, model outputs, and downstream methodological summaries. The workflow was organised as modular steps with explicit input and output files, allowing each stage to be rerun, inspected, and validated independently.

##### S8.1 Software environment

The main software tools and package versions used in the final workflow runs are listed in Table **S25**. Version numbers correspond to the execution environment used for the final workflow runs.

**Supplementary Table S25**: Software versions used in the computational workflow.

| Software or package | Version | Role in the workflow |
|---|---|---|
| Python | 3.11.5 | General scripting, data processing, model training, and downstream analyses |
| Snakemake | 9.19.0 | Workflow orchestration and dependency management |
| Sylphy | 0.2.0 | One-hot encoding and pretrained protein language model representation extraction |
| BioSieve | 0.1.0 | Redundancy reduction and dataset partitioning |
| MMseqs2 | 18.8cc5c | Homology-based sequence clustering |
| NumPy | 2.4.6 | Numerical array operations |
| pandas | 3.0.2 | Tabular data processing and aggregation |
| SciPy | 1.17.1 | Scientific-computing utilities |
| scikit-learn | 1.8.0 | Classical machine learning models, preprocessing, and performance metrics |
| XGBoost | 3.2.0 | Gradient-boosted tree classifier |
| Matplotlib | 3.10.8 | Figure generation and diagnostic plots |

##### S8.2 Reproducibility records and retained workflow artefacts

All downstream summaries were generated from saved workflow outputs rather than from manual recalculation. Retained artefacts included curated and harmonised datasets, representation matrices, redundancy-reduction outputs, sequence-to-representative mappings, split files, split summaries, model-output tables, metric summaries, train–test similarity diagnostics, paired candidate–reference tables, delta summaries, validity records, and figure-ready aggregation files.

The workflow metadata preserved the distinction between the representation used for model training, the representation or criterion used for redundancy control, and the representation used for distance-aware splitting. This separation allowed matched methodological comparisons to be reconstructed from saved configuration identifiers and paired-comparison tables. Invalid splits, incomplete model runs, and unmatched candidate–reference configurations were recorded and excluded from aggregate summaries according to the validity criteria described in Section S6.

The main artefacts retained to support reproducibility are summarised in Table **S26**. These records allow the full workflow to be audited from source-level dataset harmonisation to downstream methodological summaries.

**Supplementary Table S26**: Workflow artefacts retained to support reproducibility and reconstruction of downstream analyses.

| **Workflow stage** | **Retained artefacts and reproducibility purpose** |
|---|---|
| Dataset harmonisation | Source-level processed tables, metadata files, harmonised datasets, conflict records, and filtered final datasets were retained to preserve source provenance, label assignment, conflict handling, and sequence-level filtering decisions. |
| Representation extraction | Representation matrices, sequence identifiers, labels, and representation metadata were retained to preserve the correspondence between each protein sequence and its numerical representation. |
| Redundancy reduction | Reduced datasets, retained and removed sequence summaries, representative mappings, threshold metadata, and reduction reports were retained to document which sequences were kept or removed under each homology-based or representation-distance-based criterion. |
| Dataset partitioning | Train, validation, and test files, split parameter files, split summaries, fold reports, and invalid-split records were retained to verify split integrity, class diversity, train–test separation, and seed-specific partitions. |
| Model training and evaluation | Model-output tables, hyperparameter configuration identifiers, metric summaries, seed identifiers, and split metadata were retained to reconstruct model evaluation across representations, reductions, partitioning strategies, classifiers, and random seeds. |
| Downstream methodological analyses | Train–test similarity summaries, paired candidate–reference tables, delta summaries, performance-retention summaries, validity records, and figure-ready aggregation files were retained to reconstruct methodological comparisons and diagnostic analyses. |

Configuration files, workflow rules, execution parameters, curated datasets, processed outputs, and analysis scripts are provided with the public repository and archived outputs described in the Code and Data Availability Statement.

#### References

Ashfaq Ahmad, Shahid Akbar, Maqsood Hayat, Farman Ali, Salman Khan, and Mohammad Sohail. Identification of antioxidant proteins using a discriminative intelligent model of k-space amino acid pairs based descriptors incorporating with ensemble feature selection. *Biocybernetics and Biomedical Engineering*, 42 (2):727–735, 2022.

Saeed Ahmed, Muhammad Arif, Muhammad Kabir, Khaistah Khan, and Yaser Daanial Khan. Predaodp: accurate identification of antioxidant proteins by fusing different descriptors based on evolutionary information with support vector machine. *Chemometrics and Intelligent Laboratory Systems*, 228:104623, 2022.

Judith Bernett, David B Blumenthal, and Markus List. Cracking the black box of deep sequence-based protein–protein interaction prediction. *Briefings in Bioinformatics*, 25(2):bbae076, 2024.

Ahmad Hassan Butt, Nouman Rasool, and Yaser Daanial Khan. Prediction of antioxidant proteins by incorporating statistical moments based features into chou's pseaac. *Journal of theoretical biology*, 473: 1–8, 2019.

Salvatore Candido, Thomas Hayes, Alexander Derry, Roshan Rao, Zeming Lin, Robert Verkuil, Bryan Wu, Jin Sub Lee, Elise S. Bruguera, Jehan A. Keval, Mykhailo Kopylov, John E. Pak, Wesley Wu, Neil Thomas, Samson Mataraso, Alvin Hsu, Ashton C. Trotman-Grant, Kilian Fatras, Allan dos Santos Costa, Rohil Badkundri, Halil Akın, Deniz Oktay, Jonathan Deaton, Elizabeth Montabana, Hrishita Sitwala, Yue Yu, Marius Wiggert, Dylan Alexander Carlin, Anthony W. Goering, Tomasz Blazejewski, McCullen Sandora, Michael Hla, Tina Z. Jia, Leon H. Kloker, Nicholas J. Sofroniew, Masatoshi Uehara, Jassi Pannu, Sharrol Bachas, Daniel S. Liu, Tom Sercu, and Alexander Rives. Language modeling materializes a world model of protein biology, 2026. URL `https://www.biorxiv.org/content/10.64898/2026.06.03.729735`. Preprint.

Ahmed Elnaggar, Michael Heinzinger, Christian Dallago, Ghalia Rehawi, Yu Wang, Llion Jones, Tom Gibbs, Tamas Feher, Christoph Angerer, Martin Steinegger, et al. Prottrans: toward understanding the language of life through self-supervised learning. *IEEE transactions on pattern analysis and machine intelligence*, 44(10):7112–7127, 2021.

Ahmed Elnaggar, Hazem Essam, Wafaa Salah-Eldin, Walid Moustafa, Mohamed Elkerdawy, Charlotte Rochereau, and Burkhard Rost. Ankh: Optimized protein language model unlocks general-purpose modelling. *arXiv preprint arXiv:2301.06568*, 2023.

Peng-Mian Feng, Hao Lin, and Wei Chen. Identification of antioxidants from sequence information using naive bayes. *Computational and mathematical methods in medicine*, 2013(1):567529, 2013.

Pengmian Feng, Hui Ding, Hao Lin, and Wei Chen. Aod: the antioxidant protein database. *Scientific reports*, 7(1):7449, 2017.

Enrique Fernández-Blanco, Vanessa Aguiar-Pulido, Cristian Robert Munteanu, and Julian Dorado. Random forest classification based on star graph topological indices for antioxidant proteins. *Journal of theoretical biology*, 317:331–337, 2013.

Luu Ho Thanh Lam, Ngoc Hoang Le, Le Van Tuan, Ho Tran Ban, Truong Nguyen Khanh Hung, Ngan Thi Kim Nguyen, Luong Huu Dang, and Nguyen Quoc Khanh Le. Machine learning model for identifying antioxidant proteins using features calculated from primary sequences. *Biology*, 9(10):325, 2020.

Xiaoyang Jing, Qiwen Dong, Daocheng Hong, and Ruqian Lu. Amino acid encoding methods for protein sequences: a comprehensive review and assessment. *IEEE/ACM transactions on computational biology and bioinformatics*, 17(6):1918–1931, 2019.

Rujun Li, Haotian Wang, Qiunan Yu, Jing Cai, Liangzhen Jiang, Ximei Luo, Quan Zou, and Zhibin Lv. Aopxsvm: A support vector machine for identifying antioxidant peptides using a block substitution matrix and amino acid composition, transformation, and distribution embeddings. *Foods*, 14(12):2014, 2025.

Zeming Lin, Halil Akin, Roshan Rao, Brian Hie, Zhongkai Zhu, Wenting Lu, Nikita Smetanin, Allan dos Santos Costa, Maryam Fazel-Zarandi, Tom Sercu, Sal Candido, et al. Language models of protein sequences at the scale of evolution enable accurate structure prediction. *bioRxiv*, 2022.

David Medina-Ortiz, Diego Álvarez, and Julián García-Vinuesa. Sylphy: Protein sequence representation: Encoders, embeddings, and reductions. `https://pypi.org/project/sylphy/`, 2026a. Python package, version 0.2.0, accessed 12 May 2026.

David Medina-Ortiz, Diego Álvarez-Saravia, and Julián García-Vinuesa. BioSieve: Redundancy reduction and leakage-aware dataset partitioning for biological machine learning. https://pypi.org/project/biosieve/, 2026b. Python package, version 0.1.0, accessed 12 May 2026.

Rajat Kumar Mondal, Debarup Sen, Ankish Arya, and Sintu Kumar Samanta. Developing anti-microbial peptide database version 1 to provide comprehensive and exhaustive resource of manually curated amps. *Scientific Reports*, 13(1):17843, 2023.

Raphaël Mourad. Mistral-prot-v1-134m (mistral for protein). https://huggingface.co/RaphaelMourad/Mistral-Prot-v1-134M, 2024. Hugging Face Model Repository.

Konstantin Schütze, Michael Heinzinger, Martin Steinegger, and Burkhard Rost. Nearest neighbor search on embeddings rapidly identifies distant protein relations. *Frontiers in Bioinformatics*, 2:1033775, 2022.

Martin Steinegger and Johannes Söding. Mmseqs2 enables sensitive protein sequence searching for the analysis of massive data sets. *Nature biotechnology*, 35(11):1026–1028, 2017.

Deke Sun, Ze Liu, Xiuli Mao, Zongru Yang, Chengcheng Ji, Yanxin Liu, and Shaokun Wang. Anox: a robust computational model for predicting the antioxidant proteins based on multiple features. *Analytical Biochemistry*, 631:114257, 2021.

Felix Teufel, Magnús Halldór Gíslason, José Juan Almagro Armenteros, Alexander Rosenberg Johansen, Ole Winther, and Henrik Nielsen. Graphpart: homology partitioning for biological sequence analysis. *NAR genomics and bioinformatics*, 5(4):lqad088, 2023.

Yixiao Zhai, Yu Chen, Zhixia Teng, and Yuming Zhao. Identifying antioxidant proteins by using amino acid composition and protein-protein interactions. *Frontiers in cell and developmental biology*, 8:591487, 2020.

Lina Zhang, Chengjin Zhang, Rui Gao, Runtao Yang, and Qing Song. Sequence based prediction of antioxidant proteins using a classifier selection strategy. *Plos one*, 11(9):e0163274, 2016.
